# Epistatic interaction between ribosome-associated Era GTPase and stringent response regulator RelA modulates bacterial cell growth and dormancy

**DOI:** 10.64898/2026.04.21.719802

**Authors:** Rohan Pal, Himanshu Sharma, Pallavi Sabharwal, Kutti R. Vinothkumar, Baskaran Anand

**Affiliations:** Department of Biosciences and Bioengineering, Indian Institute of Technology Guwahati, Guwahati 781039, Assam, INDIA; UCIBIO – Applied Molecular Biosciences Unit, Department of Chemistry, NOVA School of Science and Technology, NOVA University Lisbon, Caparica, Portugal Laboratory i4HB - Institute for Health and Bioeconomy, NOVA School of Science and Technology, NOVA University Lisbon, Caparica, Portugal; National Centre for Biological Sciences, Tata Institute of Fundamental Research, GKVK Post, Bengaluru, Karnataka, 560065, INDIA

## Abstract

Cellular homeostasis is sustained by balancing the cell growth and quiescence in response to specific environmental cues. While the mechanisms governing the transition from growth to dormancy are reasonably well understood, the processes underlying exit from dormancy remain poorly characterised. Using an integrated approach employing bacterial genetics, Cryo-EM, and molecular microbiology, we investigated this phenomenon by studying the cross-talk between two antagonistic pathways, ribosome biogenesis that promotes cell growth, and stringent response that induces dormancy. We show that the exit from dormancy requires a conserved Era GTPase whose synthesis selectively rises during onset of exponential growth phase in *Escherichia coli*. Era not only accelerates the maturation of head and platform regions of 30S ribosomal subunit but also dislodges RelA from 70S ribosome. This derepresses the 70S ribosome from RelA-mediated inhibition, and promotes protein synthesis driving the active cell growth. This study uncovers that temporal epistatic interaction between Era and RelA governs cellular resuscitation in bacteria. Given that the dormant bacterial populations contribute to antibiotic tolerance, understanding this regulatory axis offers new insights for resensitising the recalcitrant dormant bacteria to antibiotics.

## Introduction

A bacterial cell needs to adapt rapidly to ever-changing environmental conditions for its continued survival. In response to changes in nutrient availability or stress conditions, a bacterial cell coordinates its various cellular processes to maintain homeostasis between growth and dormancy. In bacteria, ribosome biogenesis and stringent response are two critical, mutually exclusive pathways that regulate cell growth and dormancy, respectively, ensuring bacterial adaptation and survival. During the availability of ample nutrients, bacteria synthesise new ribosomes, which correlates with high protein turnover to drive exponential growth (1, 2). Active ribosome biogenesis is the hallmark of exponential growth, and it is a highly energy-intensive process. In prokaryotes, ribosome biogenesis involves the coordinated synthesis of three different rRNAs (16S, 23S, and 5S) and more than 50 ribosomal proteins, along with numerous ribosome assembly factors (RAFs) that guide the assembly of ribosomes (3–5). For a bacterium like *Escherichia coli*, which contains ∼70,000 ribosomes to support its high growth rate (doubling time of ∼20 minutes in rich media under lab conditions), ribosome biogenesis and the protein synthesis machinery become crucial checkpoints for bacterial adaptation and essential targets for regulation by the stringent response (2, 6). Once the environment is depleted of essential nutrients or subjected to unfavourable conditions, the stringent response pathway is activated, downregulating all energy-intensive growth-associated processes, such as ribosome biogenesis and protein synthesis driving the cell towards quiescence (6, 7). A stringent response pathway regulates ribosome biogenesis in a multipronged manner by acting through its effector alarmone molecule, ((p)ppGpp), synthesised by RelA-SpoT homolog (RSH) family enzymes such as RelA in *Escherichia coli* (8, 9). Since rRNA synthesis is the rate-limiting step of ribosome biogenesis, alarmones downregulate the transcription of rRNA operons (8, 9). Alarmones also block ribosome maturation by inhibiting the activity of various ribosome associated GTPases (RA-GTPases) involved in assembly and translation (10–12).

Further alarmones promote ribosome hibernation to preserve the ribosomes from degradation during the stationary phase or stressful conditions (13). This metabolic reprogramming alters the cellular expression profile by reallocating cellular resources away from growth towards cellular maintenance of essential functions and stress resistance.

The induction of stringent response under stress shows how the cell tips the balance of growth towards dormancy. However, the regulatory mechanism underlying how cellular resuscitation occurs after quiescence is not clearly understood. Intrigued by this, we asked whether any of the assembly factors involved in ribosome biogenesis play a regulatory role in reviving growth during nutrient availability by counteracting the effects of the stringent response on the translation machinery. Towards this, we probed the role of Era GTPase, a conserved RA-GTPase involved in the maturation of the 30S ribosomal subunit (14–19). Beyond ribosome maturation, Era has been implicated in cell cycle regulation and chromosomal partitioning (14, 20, 21), thus suggesting that it is an attractive candidate to counteract stringent response. Using a tuneable protein depletion system, we demonstrate that Era-depleted *E. coli* cells enter a non-dividing, yet metabolically active dormant state reminiscent of stringent response-induced quiescence. We further show that Era counteracts RelA in a temporal manner that downregulates stringent response leading to activation of ribosome synthesis and revival of cell growth.

## Materials and methods

### Creation of Bacterial Strains

*E. coli* strain BW25113 was used as the WT parental strain for all the genetic manipulation and reference measurements. To generate the strain (K12*era*:HAssrA) for targeted depletion of essential gene *era,* we introduced *Mesoplasma florum ssr*A degradation tag at the 3’ end of the genomic copy of *era*. A Knock-in DNA cassette was prepared by amplifying the chloramphenicol acetyltransferase (CAT) cassette flanked by *loxP* site from pKD32 plasmid (22). DNA sequence encoding for HA tag followed by an optimised mf-SsrA tag was further fused at the 5’ end using overlap extension PCR. Finally, the left and right flanking regions, homologous to 50 bp upstream and downstream of the 3’ region of the *era* gene, were added to the cassette by PCR. The cassette was inserted into the chromosome using the λ-Red recombineering method with the help of pKD46 plasmid (22). Upon confirmation of integration, the CAT gene was cured by overexpressing Cre recombinase from a heat-curable plasmid.

A similar strategy was used to incorporate the HA tag at the 3’ end of the chromosomal copy of *spoT* by using the λ-Red recombineering method for detecting SpoT in *E. coli* cells. Here, the aminoglycoside phosphotransferases (KanR) gene flanked by FRT site amplified from pKD13 plasmid was used to synthesise the knock-in cassette (22). The insertion was verified by PCR and sequencing.

*relA* null mutant was prepared against the K12era:HAssrA background using Aminoglycoside adenylyltransferase (SmR) deletion cassette amplified from pET-13S-R plasmid (Addgene #48328). Deletion of the chromosomal copy of *rel*A was undertaken by λ-Red recombineering method using pKD46 plasmid and verified by PCR and sequencing (22).

### Cloning of genes

The gene encoding Era was amplified from *E. coli* genomic DNA and cloned into a pET-1R vector (Addgene #29664) using the restriction site SspI under a T7 promoter with Strep-Tag II on the N-terminus using Gibson assembly. Similarly, cloning of G-Domain (1-185 aa), KH-Domain (186-301aa), K21A mutant, ObgE and HflX were performed as described above. The gene encoding RelA was amplified from *E. coli* genomic DNA and cloned into pLJSRSF7 vector (Addgene #64693) under a T7 promoter with a 6His_MBP_SUMO tag on the N-terminus.

For *in vivo* translation assays, the reporter gene mEGFP was amplified from 1GFP plasmid (Addgene # 29663) and cloned using Gibson assembly in pQE2 using the restriction enzyme NdeI/ HindIII. Following the cloning in pQE2 vector, the mEGFP gene under the T5 promoter was further subcloned into the pET-1R vector backbone to change the antibiotic resistance to KanR by using the restriction sites XhoI/NheI and this places the mEGFP gene under the T5 promoter in pET-1R. Similarly, the β-Galactosidase gene was moved from the pQE2 vector (23) to 1R with T5 promoter.

### Purification of proteins

Era (1-301 aa) and its variants such as G-domain (1-185 aa), KH-Domain (186-301 aa), and K21A mutant, ObgE and HflX were all purified using a similar purification strategy. All the genes coding for the respective proteins were cloned in pET-1R vector with a strep-tag II on the N-terminus and expressed in *E. coli* BL21 (DE3) cells. For overexpression, the *E. coli* cells harbouring the expression plasmid were grown in 2L of LB media supplemented with 50 µg/ml Kanamycin at 37⁰C, 180 rpm. The cells were grown till OD_600_ was 0.6, and then the expression was induced using 0.3 mM IPTG and grown at 25⁰C for 16 hr. Following this, cells were harvested by centrifugation at 8000 x g, 4⁰C/10 min. Cells were washed with buffer PG1 (20 mM Tris-Cl pH 8, 500 mM NaCl, 5% glycerol, 6 mM β-ME) and resuspended in 30 ml of buffer PG1 with 1 mM PMSF and lysed by sonication. After removing the cell debris by centrifugation at 20000 x g, 4⁰C/1 hr, the clarified lysate containing the protein of interest was then bound to a StrepTrap^TM^ HP column (Cytiva) equilibrated with buffer PG1. Following the binding, the column was washed with 5 CV of buffer PG1 and eluted with 4 CV of buffer PG1 containing 2.5 mM Desthiobiotin. Eluted fractions were analysed by SDS-PAGE, and peak fractions were loaded onto a HiLoad Superdex 75 pg gel filtration column (Cytiva) equilibrated with buffer PG2 (20 mM Tris-Cl pH 8, 200 mM NaCl, 100 mM KCl, 5% glycerol, 6 mM β-ME). The peak fractions from gel filtration chromatography were analysed using SDS PAGE for homogeneity and then concentrated through a 10 kDa membrane filters (Sartorius) using ultrafiltration. The purified proteins were aliquoted in small volumes, flash frozen and stored at -80⁰C, until further use.

*E. coli* Bl21(DE3) transformed with pLJ_RelA was grown in 2L of LB supplemented with 50 µg/ml Kanamycin till OD_600_ 0.6. The expression was induced with 0.3 mM IPTG and grown for 4 hr at 37⁰C, 180 rpm. Cells were harvested at 8000 x g, 4⁰C for 10 min and washed once with buffer PR1 (20 mM Tris-Cl pH 7.5, 500 mM KCl, 1mM Mg(CH₃COO)₂, 20 mM Imidazole, 10% glycerol, 6 mM β-ME). The cells were resuspended in buffer PR1 with 1mM PMSF and lysed by sonication. The cell lysate was clarified by centrifugation at 20000 x g, 4⁰C/1hr and then loaded onto a HisTrap™ HP column (Cytiva) equilibrated with buffer PR1. The column was then washed with 5 CV of buffer PR1, and elution was carried out using a linear gradient of PR2 (20 mM Tris-Cl pH 7.5, 500 mM KCl, 1 mM Mg(CH₃COO)₂, 500 mM Imidazole, 10% glycerol, 6 mM β-ME) on an Akta Pure system (Cytiva). The elution fractions were analysed by SDS-PAGE, and peaks corresponding to pure fractions of RelA were collected and treated with an in-house purified Ulp1 protease to remove His-MBP-SUMO tag by overnight incubation at 4⁰C to yield a tag-free RelA. The cleaved tag was removed from the protein mixture by passing it through the MBPTrap™ HP column (Cytiva). The flow-through solution was passed through HiLoad Superdex 200 pg gel filtration column (Cytiva) equilibrated with buffer PR3 (20 mM Tris-Cl pH 7.5, 500 mM KCl, 10 mM NH_4_Cl, 10 mM Mg(CH₃COO)₂, 10% glycerol, 6 mM β-ME) to remove additional contaminants. The peak fractions were analysed by SDS and the purified RelA was concentrated through a 10 kDa membrane filters (Sartorius) using ultrafiltration and stored at -80⁰C in small aliquots after flash freezing.

### Depletion of Era and Growth Analysis

For tuneable and targeted depletion of Era, *E. coli* K12*era*:HAssrA cells were transformed with pBAD33-mf-lon (Addgene #21867) plasmid, where the gene coding for Lon protease from *M. florum* was expressed under a pBAD promoter. Following transformation, the cells were grown on an LB agar plate supplemented with 25 µg/ml chloramphenicol and 0.2 % glucose (to prevent leaky expression) at 37 ⁰C. Single colony was picked and incubated in a 5ml LB tube with 25 µg/ml chloramphenicol (Cmp) and 0.2 % glucose and grown at 37 ⁰C, 180 rpm. The cells were first harvested and washed with LB media to remove glucose. Following this, cells were diluted 100 times with fresh LB media supplemented with 25 µg/ml Cmp and grown in the presence or absence of 13 mM arabinose for induction of *Mf*-Lon protease at 37 ⁰C, 180 rpm. Once the cells reached the early log phase (OD_600_ ≤ 0.2), they were diluted 10-fold with fresh media supplemented with Cmp and arabinose. The procedure was repeated for one more cycle, with close monitoring of OD_600_ in a 24-well assay plate using a Tecan infinite M200 multimode plate reader. The growth profile for Era depletion against Δ*relA* background was probed similarly.

### Fluorescence microscopy

WT and Era^-^ cells were grown according to the protocol described above until the Era^-^cells were growth arrested in cycle III. The cells were then harvested and divided into three equal fractions of 200 µl. The fractions were washed with Phosphate buffer saline (PBS) by centrifugation and resuspended in 200 µl of PBS. The first two fractions with live cells were stained with 1 µg/ml acridine orange (AC) and propidium iodide (PI), respectively. The cells were incubated for 1 min in tumbling motions, and excess dye was removed by centrifugation and washing with PBS. The third fraction of bacterial cells was processed as a control by fixing with chilled methanol and acetone solution in a 1:1 ratio. It is added in a drop-wise manner over 1 min with regular inverted mixing on ice until the total volume reaches 1 ml. The cells were washed with PBS and stained with 1 µg/ml propidium iodide. Excess dye was removed by centrifugation and washing with PBS. 5 µl was added to an agarose pad, covered with a coverslip and visualised under oil immersion-100X objective using an Olympus CKX53 microscope with a CoolLED pE-300^white^ fluorescence light source. Acridine orange was analysed by blue excitation/green emission fluorescence mirror unit. Propidium iodide was analysed using the green excitation/red emission fluorescence mirror unit. The images were captured using cellSens software (Olympus) and further processed using Fiji (24) to calculate the cell length.

### MTT assay

WT and Era^-^ cells were grown according to the aforementioned depletion protocol till cycle III. Cells were harvested and normalised to OD_600_ 1.0 for both WT and Era^-^ in LB media. Also, one set each of WT and Era^-^ was heat treated (HT) at 95⁰C for 10 min to be used as control. 200 µl each of the sample-only media (Blank), WT, WT (HT), Era^-^, Era^-^ (HT) was taken, and 50 µl of MTT (3-(4,5-dimethylthiazol-2-yl)-2,5 diphenyl tetrazolium bromide)-solution (5 mg/ml) was added to each sample and incubated at 37⁰C for 3 hr. 50 µL of DMSO was added to solubilise the formazan crystals and incubated for 15 min after rigorous vortexing. The sample was centrifuged to remove the cell pellet, 200 µl of the upper supernatant was added to 96 well assay plate, and absorbance was recorded at 570 nm using a Tecan infinite M200 multimode plate reader.

### *in vivo* translation assay

WT and Era^-^ cell were co-transformed with the reporter plasmid pT5_mEGFP and grown to Era depleted state. The reporter gene was induced using 0.1 mM IPTG and grown for a further 30 min at 37 ⁰C, 180 rpm. 200 µl cells were harvested by centrifugation and washed with PBS. Cells were resuspended in 50 µl PBS. 5 µl of the sample was loaded onto 1.5% agarose pads with a coverslip. The samples were immediately visualised under oil immersion-100X objective using the fluorescence microscopy system as described previously. The fluorescence intensity of the excitation light source was set to 4, and images were captured at an exposure time of 0.158 s for all samples using cellSens software (Olympus). The images were further processed in Fiji (24) and MicrobeJ (25). The area of each bacterial cell and the respective fluorescence intensity in each cell were measured. The fluorescence units were normalised with respect to the cell area as follows :

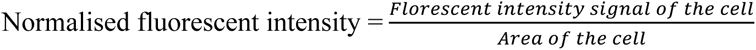

### Reverse transcription PCR

1 ml cell samples from the previous *in vivo* translation assay were processed to isolate total cellular RNA by Trizol-mediated extraction (Thermofisher). The total RNA sample was treated with DNase I (NEB) and the sample was purified through a second phenol-chloroform-mediated extraction and ethanol precipitation. 50 ng of total RNA from each sample was processed for first-strand cDNA synthesis by SuperScript ™ III RT (Invitrogen) using a primer specific to 3’ end of the mEGFP gene. The cDNA product was then used as a template for PCR amplification using gene specific primers. The amplified sample was run on a 1% TAE agarose gel and visualised by EtBr staining using the Bio-Rad ChemiDoc^TM^ XRS+ gel documentation system.

### Polysome profiling

WT and Era^-^ cells were grown according to the protocol described above until the Era^-^cells were growth arrested in cycle III (OD_600_ ≤ 0.2). The depletion was slightly modified to allow the process to be upscaled to 2L culture. At this point, the culture was chilled rapidly on ice with the addition of ice cubes and 100 µg/ml Cmp, followed by incubation for 15 min. The cells were harvested by centrifugation at 4 °C, 8000 × g for 10 min. Cells were then washed once with Buffer RL (20 mM Tris–HCl (pH 7.5 at 25°C), 150 mM NH_4_Cl, 10.5 mM Mg(CH₃COO)₂, 0.5 mM EDTA and 6 mM β-ME) and resuspended in 1 ml buffer RL supplemented with 1 mg/ml Lysozyme, 1 mM PMSF, 0.05× CellLyticB (Sigma) and 10 U of DNase I (NEB) followed by incubation on ice for 1 h. The cells were lysed by five cycles of freeze-thaw using liquid nitrogen. The lysate was clarified by centrifugation at 20000 x g, 4 ⁰C, 1 hr. The cleared lysate was transferred to a fresh tube, and the absorbance at 260 nm was quantified using an Implen Nanospectrophotometer. 100 µl of 10000 A_260_ lysate was loaded onto a 10%-50% sucrose gradient in buffer RL and ultracentrifuged at 78600 g, 4 ⁰C, 15 hr in a TH-641 (Thermo-Sorvall WX+ ultracentrifuge) rotor for separation of the ribosomes. The fractionation and profiling was performed using a syringe pump connected to the Akta Pure system (Cytiva).

To perform polysome profiling from the lag phase, K12era:HAssrA cells were grown overnight in a 5 ml LB media at 37 ⁰C at 180 rpm. 1% inoculum was transferred to 1.5 L LB media in a 5 L flask and grown at 37 ⁰C with shaking at 180 rpm. When the cells reached an OD_600_ of 0.05, the cells were immediately chilled with ice cubes in the presence of freshly prepared 1.1% (final concentration) formaldehyde and incubated for 12 min on ice. The cross-linking was stopped by adding 250 mM (final concentration) glycine. The cells were harvested by centrifugation at 8000 x g for 10 min at 4 ⁰C. The cells were washed with buffer CL (20 mM HEPES-KOH (pH 7.4 at 25 °C), 100 mM KCl, 10.5 mM Mg(CH₃COO)₂ and 0.5 mM EDTA). Cells were lysed and profiled similar to polysome profiling mentioned earlier. The sample was fractionated (500 µL fractions) and concentrated by TCA precipitation, and the pellet was processed by resuspending in 1X SDS-PAGE dye. The fractions were resolved on a 12% SDS-PAGE. The presence of Era and RelA in the fractions was probed by 1:10000 mouse anti-HA Tag monoclonal primary antibody (Invitrogen) and 1:1000 mouse anti-RelA monoclonal primary antibody (Santa Cruz Biotechnology), respectively. They were then probed using 1:5000 goat anti-mouse HRP conjugated secondary antibody (Invitrogen).

### Purification of Ribosomal particles

To purify 70S ribosomal particles, WT cells were grown overnight in LB media. 1% of the inoculum was added to 400 ml LB and grown with shaking at 180 rpm till OD_600_ reaches 0.6. The cells were rapidly harvested by centrifugation at 8000 x g, 4 ⁰C for 10 min. Cells were then washed once with Buffer RL (20 mM Tris–HCl (pH 7.5 at 25 °C), 150 mM NH_4_Cl, 10.5 mM Mg(CH₃COO)₂, 0.5 mM EDTA and 6 mM β-ME) and later resuspended in 1 ml buffer RL supplemented with 1 mg/ml Lysozyme, 1 mM PMSF, 0.05× CellLyticB, 10 U of DNase I reagent followed by incubation on ice for 1 h. The cells were lysed by five cycles of freeze-thaw using liquid nitrogen. The lysate was cleared of cellular debris by centrifugation at 20000 x g, 4 ⁰C, 1 hr. The lysate was made up to 5 ml using buffer RL and layered onto a 6 ml sucrose cushion composed of buffer RC (20 mM Tris–HCl (pH 7.5 at 25°C), 1 M NH_4_Cl, 10.5 mM Mg(CH₃COO)₂, 0.5 mM EDTA, 1.1 M sucrose and 6 mM β-ME) and centrifuged at 100000 x g, 4 ⁰C for 18 hr on TH-641 rotor. The upper supernatant was discarded, and the crude ribosomal pellet was first washed with buffer RL and followed by resuspension in 600 µL of buffer RL supplemented with 60 U of SUPERase·In™ RNase Inhibitor slowly over 4 hr on a rocker. Absorbance at 260 nm was quantified for the ribosomal sample by Implen Nanospectrophotometer, and a total of 100 µl of A_260_=10000 was loaded onto a 10%-50% sucrose gradient in buffer RL and ultracentrifuged at 78000 g, 4 ⁰C, 15 hr in a TH-641 rotor for separation of the ribosomes. The fractionation was performed using a syringe pump connected to the Akta Pure system (Cytiva). 70S fractions were collected and concentrated by ultracentrifugation at 100000 x g, 4 ⁰C, 18 hr using TH-641 rotor. The 70S ribosomal pellet was then resuspended in buffer RS (20 mM Tris–HCl (pH 7.5 at 25 °C), 60 mM NH_4_Cl, 10 mM Mg(CH₃COO)₂ and 4 mM β-ME) and stored at -80 ⁰C in small aliquots by flash freezing.

For the isolation of mature 30S and 50S ribosomal subunits, the step followed was similar to the above-mentioned protocol with slight adjustments. After pelleting the crude ribosomes using sucrose cushion, the crude ribosomes were dissociated in buffer RD (20 mM Tris–HCl (pH 7.5 at 25 °C), 30 mM NH_4_Cl, 1 mM Mg Mg(CH₃COO)₂, 0.1 mM spermidine and 6 mM β-ME) for 2 hr with gentle rocking on ice and immediately loaded onto a 10-35% sucrose gradient in buffer RL and ultracentrifuged at 50000 g, 4 ⁰C, 15 hr for fractionation of individual 30S and 50S subunits. The ribosomal particles thus purified were used for all the *in vitro* assays.

For purification of 30S and 70S ribosomes for Cryo-EM, Era^-^ cells were grown as described earlier until the cells are growth arrested in cycle III (OD_600_ ≤ 0.2). The depletion was slightly modified to allow the process to be upscaled to 2L of LB media. The cells were immediately harvested by centrifugation at 4 °C, 8000 × g for 10 min. The purification protocol was similar as described above. Following purification, the 30S, 50S and 70S ribosomal pellet was then resuspended in buffer CS (20 mM HEPES-KOH (pH 7.5 at 25°C), 50 mM KCl, 10 mM NH_4_Cl, 10 mM Mg(CH₃COO)₂ and 6 mM β-ME) and stored at -80 °C in small aliquots of 5 µM by flash freezing.

### *in vitro* translation assay

For *in vitro* translation assay (IVT), we used the PURExpress® Δ Ribosome Kit (NEB) and the reactions were set according to the prescribed protocol. To test the translation efficacy of ribosome particles, we purified ribosomal particles from WT and Era^-^ cells. The reaction was reconstituted with 1µM of 30S and 50S subunits from WT and Era^-^ cells, 50 ng of 1GFP plasmid and 5 U of SUPERase·In™ RNase Inhibitor (Invitrogen). Reactions were incubated at 37⁰C for 2 hr for the synthesis of the reporter gene (mEGFP). Following incubation, the reactions were diluted with 100 µl of Buffer A (20 mM Tris-Cl pH 7.5, 150 mM NaCl). The fluorescence spectrum was measured for the mEGFP reporter by excitation at 488 nm and scanned for emission from 500 to 600 nm, with averaging over three scans after baseline correction in a FluoroMax-4 spectrofluorometer (Horiba Scientific). The slit width used for excitation and emission was 2 and 5 nm, respectively.

To assess the effect of Era on translation, IVT reactions as described above were set up in the presence of Era. Purified Era was added incrementally (10-60 µM) to the individual reaction mix supplemented with WT 30S and 50S of 1 µM each. Following the incubation for 2 hr at 37⁰C, the samples were processed by heating in 1X SDS-PAGE gel loading buffer and resolved on a 12% SDS-PAGE gel followed by western blotting. The expression of the mEGFP reporter gene was probed using a 1:10000 rabbit anti-GFP primary antibody (Bio Bharati Life Science) and 1:8000 goat anti-rabbit HRP conjugated secondary antibody (Invitrogen). The blot was developed by Clarity Western ECL Substrate (Bio-RAD) and visualised using the BIO-RAD ChemiDoc^TM^ XRS+ documentation system. The signal intensity was calculated using Image lab software (Bio-RAD) from the developed blot. The IVR reaction without Era was used as a control. Relative intensity was calculated by the following formula:

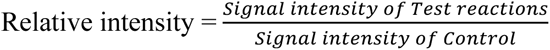

To study the effect of (p)ppGpp and Era the IVT reactions were supplemented with 2 mM ppGpp (Jena biosciences), followed by the addition of Era. To probe the effects of RelA and Era, we incubated IVT reaction containing 0.25 µM WT 70S ribosomes with 1 µM RelA, 25 µM ppGpp, and 2.5 µM Era. The assay was incubated at 37⁰C for 1 hr. The reactions were processed and analysed as described earlier.

### β Galactosidase assay

K12*era*:HAssrA and K12*era*:HAssrAΔ*relA* cells were transformed with pBAD33-mf-lon and pT5_βgal and grown till depletion of Era until Cycle III. The WT, RelA^-^, Era^-^ and RelA^-^Era^-^ cells were all normalised to OD_600_ 0.2. These cultures were then shifted to a water bath at 37°C, and β-gal production was induced with 1 mM IPTG, followed by instant mixing for 5 s. 200 µl sample was drawn from the induced cultures every 30 s till 540 s and added to MCT tubes containing 300 µl Z-buffer (60 mM Na_2_HPO_4_·H_2_O, 40 mM NaH_2_PO_4_·H_2_O (pH 7.0 at 25°C), 10 mM KCl, 1 mM MgCl_2_, 5 mM β-ME), 20 μl chloroform, and 10 μl 0.1% SDS. The tubes were vortexed for 10 s and transferred to ice. All the tubes were incubated for 5 min on ice and then for 5 min at RT, after which 500 μl of 1 mg/ml o-nitro-phenyl galactopyranoside (ONPG) was added to the reaction. ONPG hydrolysis was allowed to proceed for 5 min at RT to develop a chromogenic product. The reaction was stopped after adding 500 μl of 1 M Na_2_CO_3_. The colour was allowed to be stabilised for 10 min, following which the tubes were centrifuged at 13000 x g, 1 min and 300 µl of the reaction mix was transferred to 96-well plates. Absorbance was recorded at 420 and 550 nm using a Tecan infinite M200 multimode plate reader.

### Quantitative real-time PCR

To probe the expression of *era*, *relA* and *spoT* in bacterial cells, *E. coli* K12 cells (WT) were grown in LB media. Samples were harvested at the following time point (hr): 0, 0.5, 1, 1.5, 2, 2.5, 3, 4, 6 and 8, covering lag, log and stationary phases of the growth. The cells were harvested by centrifugation at 13000 x g for 2 min. The cells were washed with buffer A (20 mM Tris-Cl pH 7.5, 150 mM NaCl) and resuspended in sodium citrate buffer 5.2 pH. The total cellular RNA was isolated from each time point samples by hot acid phenol extraction followed by silica column based isolation. The sample was treated with DNase I (NEB) to remove DNA contamination. The total RNA samples were subjected to a sample clean-up step using silica column based method. Total RNA concentration was quantified using Qubit RNA BR Assay kit (Invitrogen). A total of 2 µg total RNA was taken for each time point sample and reverse transcription was carried to synthesise cDNA using SuperScipt^TM^ III Reverse Transcriptase (Invitrogen) using 1X RT random primers (Applied Biosystems) as per the kit described protocol. The cDNA reactions were diluted using elution buffer and a total of 10 ng was used as a template for quatitative real-time PCR using the iTaq Universal SYBR Green Supermix (Bio-RAD) (10 µl per reaction) with 500 nM forward and reverse primers each specific to *era*, *rel*A, *spo*T, *gyr*A, and *cys*G across different time points. Real-time PCR was performed using AriaMax Real-Time PCR thermocycler (Agilent) in 8-tube clear PCR strips (Bio-RAD) sealed with Optical Flat 8-Cap strips (Bio-RAD). The RT-qPCR temperature profile included an initial denaturation step at 95 °C for 10 min, followed by 36 cycles of 15 s at 95 °C, and 15 s at 60 °C. A melting curve was performed at the end of the RT-qPCR run by stepwise (0.5 °C per 5 s) increasing the temperature from 60 °C to 95 °C. All experiments were carried out in three independent biological replicates. For each cDNA batch, two independent RT-qPCR runs were performed for each gene. The Cq value was determined by using the same baseline threshold for all experiments using the Agilent AriaMx software. The geometric mean of Cq values of two reference gene *gyrA* and *cysG* was used for normalisation to calculate the ΔCq value across the time points for *era*, *relA* and *spoT*. Relative expression for the respective gene was calculated as 2^-ΔCq^.

### Protein profiling

To profile the synthesis of Era, RelA, and SpoT proteins, *E. coli* K12*era*:HAssrA and K12*spoT*:HAssrA were grown in LB media. Samples were harvested at the following time point (hr): 0.5, 1, 1.5, 2, 2.5, 3, 4, 6 and 8, covering lag, log and stationary phases of the growth. The cells were harvested by centrifugation at 8000 x g for 10 min. The cells were washed with buffer A (20 mM Tris-Cl pH 7.5, 150 mM NaCl) and resuspended in buffer A. The cells were lysed by sonication, and the lysate was cleared by centrifugation. Total protein concentration was quantified by Qubit Protein Broad Range Assay (Invitrogen) for lysate associated with each time point, and the total protein concentration in all samples was normalised. The cell lysate was processed by heating with 1X SDS-PAGE dye, and 25 µg of total protein was loaded per sample point and resolved on a 12% SDS-PAGE gel followed by western blot. Era_HA and SpoT_HA were probed by 1:5000 mouse anti-HA monoclonal primary antibody (Invitrogen). RelA was probed using a 1:1000 mouse anti-RelA monoclonal primary antibody (Santa Cruz Biotechnology). All the proteins were then probed using 1:10000 goat anti-mouse HRP conjugated secondary antibody (Invitrogen). GAPDH was used as an internal loading control and probed using 1:10000 mouse anti-GAPDH monoclonal primary antibody (Invitrogen) and 1:20000 goat anti-mouse HRP conjugated secondary antibody (Invitrogen). The blot was developed using Clarity Western ECL Substrate (Bio-RAD) and visualised using the BIO-RAD ChemiDoc^TM^ XRS+ documentation system.

### *in vitro* ribosome binding assays

Ribosome co-sedimentation assay was used to determine the interaction between the 30S ribosomal subunit and Era. 1 µM 30S ribosomal subunit was incubated in the presence of an increasing concentration of Era (0 to 20 µM) with 0.5 µM GMPPNP in the reaction buffer (20 mM Tris–HCl (pH 7.5 at 25°C), 30 mM KCl, 10 mM Mg(CH₃COO)₂ and 4 mM β-ME) to a total reaction volume of 50 µl. The binding reactions were incubated for 15 min at 37⁰C in a water bath. subsequently, the reaction mixture was layered onto a 1.5 ml 15% sucrose cushion prepared in buffer RS (20 mM Tris–HCl (pH 7.5 at 25°C), 60 mM NH_4_Cl, 10 mM Mg(CH₃COO)₂ and 4 mM β-ME) to pellet the ribosomes by ultracentrifugation at 200000 x g at 4⁰C for 2 hr using the F50L-24×1.5 (Thermofisher) rotor. The 30S bound Era in the pellet was immediately separated from the unbound Era in the supernatant by transferring to a fresh MCT tube. The unbound Era in the supernatant was concentrated by TCA-mediated precipitation. The ribosomal pellet and supernatant samples were processed and resolved on 12% SDS-PAGE. The presence of Era was probed by western blotting using 1:5000 mouse anti-Strep II monoclonal primary antibody (ABclonal technology) and 1:10000 goat anti-mouse HRP conjugated secondary antibody (Invitrogen). The signal intensity of Era associated with the 30S and free Era in the supernatant was quantified by densitometry. The total signal was calculated by summing up the signal for the pellet band (30S-bound Era) and the corresponding supernatant (free Era). Using the following formula, these values were used to determine the fraction of Era co-migrating with 30S.

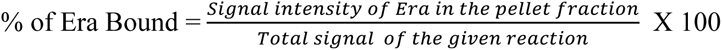

For calculating the binding affinity, the intensity corresponding to Era bound to 30S was plotted against the total amount of Era added to the reaction. The graph was then fitted using nonlinear regression (one site–specific binding) in GraphPad Prism.

### Dissociation assay

To dissociate the 70S particles to 30S and 50S subunits, 1 µM WT 70S ribosome was mixed with 10 µM of purified Era/ObgE/HflX in the presence or absence of 2mM GTP in reaction buffer (20 mM Tris–HCl (pH 7.5 at 25°C), 30 mM KCl, 5 mM Mg(CH₃COO)₂ and 4 mM β-ME) to a total volume of 20 µL. The reaction was then incubated at 37⁰C in a water bath for 30 min. Following incubation, the reaction mixture was loaded onto a 10-35% gradient cushion in buffer RS (20 mM Tris–HCl (pH 7.5 at 25°C), 60 mM NH_4_Cl, 10 mM Mg(CH₃COO)₂ and 4 mM β-ME) and ultracentrifuged at 50000 x g, 4⁰C for 15 hr using TH-641 rotor. The profiling was performed using a syringe pump connected to the Akta Pure system (Cytiva).

### Anti-association assay

To assess the association of 30S and 50S to form 70S particles, 1 µM of WT 30S was pre-incubated with 10 µM of Era in the presence or absence of nucleotides (GTP/GDP/GMPPNP) with reaction buffer (20 mM Tris–HCl (pH 7.5 at 25°C), 30 mM KCl, 20 mM Mg(CH₃COO)₂ and 4 mM β-ME) for 15 min at 37⁰C in a water bath. Following incubation, 1 µM of WT 50S was added to the reaction mix and incubated for a further 1 hr at 37⁰C. The total volume of 25 µL was then loaded onto a 10-35% gradient cushion in buffer RS (20 mM Tris–HCl (pH 7.5 at 25°C), 60 mM NH_4_Cl, 10 mM Mg(CH₃COO)₂ and 4 mM β-ME) and ultracentrifuged at 50000 x g, 4⁰C for 15 hr using TH-641 rotor. The profiling was performed using a syringe pump connected to the Akta Pure system (Cytiva).

### Pull-down assay

To assess the physical interaction between Era and RelA, we performed a pull-down assay. Magnetic Protein G beads were incubated with 2 % BSA and incubated for 1 hr at 4⁰C. Following incubation, the magnetic beads were washed 3 times with 200 µl of buffer W (20 mM Tris–HCl (pH 7.5 at 25°C), 150 mM NaCl, 30 mM KCl, 10 mM Mg(CH₃COO)₂, 4 mM β-ME and 0.2% Tween 20). The Protein G beads were then bound to 1:5000 mouse anti-Strep II-Tag monoclonal primary antibody (ABclonal technology) and incubated for 1 hr at 4⁰C. The Protein G beads were washed 3 times with 200 µl of buffer W. The Anti-Strep II-Tag antibody loaded protein G beads were mixed with 0.5 µM Era having N-terminal Strep-Tag II and incubated for 2 hr at 4⁰C in buffer B (20 mM Tris–HCl (pH 7.5 at 25°C), 150 mM NaCl, 30 mM KCl, 10 mM Mg(CH₃COO)₂ and 4 mM β-ME). The beads were washed 3 times with buffer W to remove unbound proteins. The Era-bound protein G beads were then used as a bait to pull-down varying concentrations (0 to 2 µM) of RelA by incubating in buffer B for 30 min at RT in a tumbling motion for equal mixing. The excess proteins were then washed 3 times with 200 µl of buffer W. The beads were resuspended in 25 µl of 1X SDS-PAGE dye and heated. The entire volume was loaded onto a 12% SDS-PAGE gel. The presence of both RelA and Era in pellet and supernatant fractions was analysed by western blotting. RelA was probed using a 1:1000 mouse anti-RelA monoclonal primary antibody (Santa Cruz Biotechnology), and Era was probed using a 1:5000 mouse anti-Strep II-Tag monoclonal primary antibody (ABclonal technology). This was followed by probing using 1:10000 goat anti-mouse HRP conjugated secondary antibody (Invitrogen). Varying known concentrations of RelA and Era were also simultaneously processed by western blot to be used as a standard. The signal intensity of RelA and Era was compared to the standard to quantify the concentration of the respective proteins. The pull-down index was calculated by the given formula.

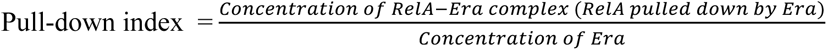

For calculating the binding affinity, the values of the Pull-down index were plotted against the total amount of RelA added to the reaction. The graph was then fitted using nonlinear regression (specific binding with Hill constant) in GraphPad Prism.

Similar procedures were followed for assessing the binding interaction of RelA with G-domain, and KH-domain of Era as well for the WT in the presence of 25 mM GMPPNP.

### RelA displacement assay

To assess the displacement of ribosome-bound RelA by Era, we devised a pelleting assay. 1µM of WT 70S ribosome was preincubated with 1µM of purified RelA along with the presence or absence of 1 µM GTP in reaction buffer (20 mM Tris–HCl (pH 7.5 at 25°C), 30 mM KCl, 10 mM Mg(CH₃COO)₂ and 4 mM β-ME) at 37⁰C for 30 min in a water bath. Following incubation, the reaction mixture was challenged with 5 µM of purified WT, G-domain, KH-domain and K21A, as the case may be, and incubated for a further 30 min at 37⁰C. Following incubation, the reactions totalling 25µl were layered onto 1.5 ml 15% sucrose cushion in buffer RS (20 mM Tris–HCl (pH 7.5 at 25°C), 60 mM NH_4_Cl, 10 mM Mg(CH₃COO)₂ and 4 mM β-ME) and centrifuged at 200000 x g for 2 hr at 4⁰C using the rotor F50L-24×1.5 rotor. The upper supernatant was immediately removed to a fresh MCT, and the free proteins were concentrated through TCA precipitation. The ribosomal pellet and the TCA precipitated supernatant were processed by 1X SDS-PAGE loading dye and resolved using a 12% SDS-PAGE. The presence of both RelA and Era in pellet and supernatant fractions was analysed using western blotting. RelA was probed using a 1:1000 mouse anti-RelA monoclonal primary antibody (Santa Cruz Biotechnology), and Era was probed using a 1:5000 mouse anti-Strep II-Tag monoclonal primary antibody (ABclonal technology). This was followed by probing using 1:10000 goat anti-mouse HRP conjugated secondary antibody (Invitrogen) for both primary antibodies. The signal intensity of RelA from the developed blot was calculated for each sample in both the pellet and supernatant fractions using densitometry. The sum of the signal in the pellet and supernatant is equivalent to the total signal of RelA for the corresponding sample. The following quantified value was used to calculate the fraction of RelA:

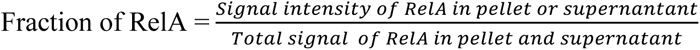

### Cryo-EM sample preparation

Initially, the ribosome sample was frozen on holey grids with a thin layer of carbon, and this gave rise to limited views of the molecule and anisotropic maps. Subsequently, 30S Era^-^ was applied to Quantifoil R 1.2/1.3 grids. The grids were glow discharged with a Pelco instrument at 25 mA for 1 minute. For the 70s ribosomes from Era^-^, the sample was applied to Quantifoil R 2/2 grids with a thin layer of carbon added. The grids were glow discharged with a Pelco instrument at 15 mA for 10 seconds. For both specimens, the Vitrobot Mark IV (ThermoFisher) was maintained at 16 °C and 100% RH, and 3 μl sample was applied to grid with a wait time of 10 seconds and followed by blotting for 3-3.5 seconds and plunge frozen in liquid ethane. The grids were stored in liquid nitrogen till imaging.

### Cryo-EM data collection and image processing

The grids were mounted on Titan Krios autogrid in liquid nitrogen and transferred to the autoloader. Grids were screened with low dose and data collection was performed with Falcon 3 detector in integration mode at a nominal magnification of 75000x, corresponding to a pixel size of 1.07 Å.

For 30S Era^-^, images were collected at a dose of ∼1.019 e^-^/Å^2^/frame and 32 frames. The movie frames were aligned using Motioncorr in RELION 4.0 (26–28) and CTF was estimated with GCTF (29). Summed images were selected using CTF values (resolution) and particles were picked with Autopick routine in RELION. Binned particles at 2.14 Å sampling were subjected to two rounds of 2D classification and good particles were selected and extracted with a box size of 320 and 1.07 Å. The particles were imported to CryoSPARC (version 4.4) (30) and one more round of 2D classification was performed. Ab-initio model generation of selected particles with 6 classes, followed by heterogenous refinement, revealed two prominent classes with well resolved features. Homogenous refinement with global and local CTF refinement were performed on these two classes with an estimated global resolution of 3 and 3.1 Å (at FSC 0.143), respectively.

For Era^-^ 70S, images were collected at a dose of ∼1.17 e^-^/Å^2^/frame and 25 frames. The movie frames were aligned using Motioncorr in RELION 4 (26–28) and CTF was estimated with GCTF (29). Summed images were selected using CTF values (resolution) and particles were picked with Autopick routine in RELION. Binned particles were subjected to two rounds of 2D classification, and good particles were selected and extracted with a box size of 360 pixels. The full data set with 746,750 particles were used to calculate an initial model and a 3D refinement was performed. This gave a map of ∼3.1 Å. These particles were subjected to 3D classification with resolution limited to 7 Å and into 6 classes. Of these, one class had only the large subunit (K=4) and one class (K=1) had low resolution feature, which was not processed further. The remaining classes and the class with only 50S subunit were subjected to 3D refinement and Bayesian Polishing and the box size was increased to 384 pixels in this step. The 3D refinement of B-factor weighted particles gave resolutions between 3.2 to 3.6 Å, and further CTFRefine was performed. To improve the quality of the map, subsequent 3D classification of each class was performed with resolution limited to 6 Å. These provided populations of empty 70S ribosomes with difference in the head domain of the 30S subunit and those with tRNA. The complete workflow of the data processing is provided in supplementary information (Fig. S2). The resolutions reported are from the postprocess step in Relion and a separate mask was used for each class. Local resolution was estimated with Relion.

All maps were analysed, and figures were prepared using UCSF ChimeraX (31). The Cryo-EM structure (PDB: 7K00) was used as a reference for analysis of all maps unless specified otherwise.

### Model Building for 30S subunit

The crystal structure of the *E. coli* ribosome (PDB: 4V4Q) was used as initial template. The mature 30S subunit was first rigid body docked in maps for Class I followed by successive rounds of real space refinement in PHENIX (32) and manual model building in Coot (version 0.91) (33). Model refinement was performed against the combined map of Class I (3.01 Å) with PHENIX using phenix.real_space_refine (version 1.14–3260) and using rRNA restraints derived from the reference model. Model validation was performed according to Brown et al (34). Areas of the map with weak or no corresponding density were deleted and not incorporated into the final model. The missing region of the model, mainly comprising the 30S head region were rigid body docked into the map for analysis.

## Results

### Depletion of Era arrests cell growth

Era, an essential GTPase in *E. coli,* is part of the *rnc* operon (35, 36) and therefore to avoid inducing any polar effects due to genetic perturbations in the operon, we chose not to introduce gene deletion but rather adopted a strategy that targets Era at the protein level. To achieve this, we employed a targeted protein degradation approach using the SsrA degron from *M. florum* (Fig. 1A) (37, 38). The chromosomal copy of *era* was fused in-frame with the *M. florum* SsrA degradation tag using Lamda Red-mediated recombineering. A plasmid-derived gene encoding *M. florum* Lon protease (mf-Lon) under an arabinose-inducible promoter was used to degrade the SsrA-tagged Era in a tuneable fashion. When Era was depleted (Era^-^) in a controlled manner using this system, we observed severe growth-associated defects (Fig. 1B). This finding is consistent with earlier studies, which have shown that perturbing Era leads to cell cycle arrest (14, 20, 21). The growth-arrested cells were stained using Acridine orange (AC) and analysed for their change in cellular morphology using fluorescence microscopy. Cells with an elongated phenotype (Fig. 1C) having an average cell length of ≥ 4 microns were observed in Era^-^ cells, indicating defects in cell division (Fig. 1D). The viability of the growth-arrested cells was further tested using propidium iodide (PI), which is impermeable to the membrane and thus stains the membrane-compromised dead cells. Era^-^ cells stained negative to PI even in a growth-arrested state, suggesting that the membrane integrity is not compromised (Fig. 1C). Using an MTT based cell viability assay, we probed the metabolic state of Era^-^ cells that suggested that Era^-^ cells are metabolically active but dormant (Fig. 1E).

**Figure 1.**
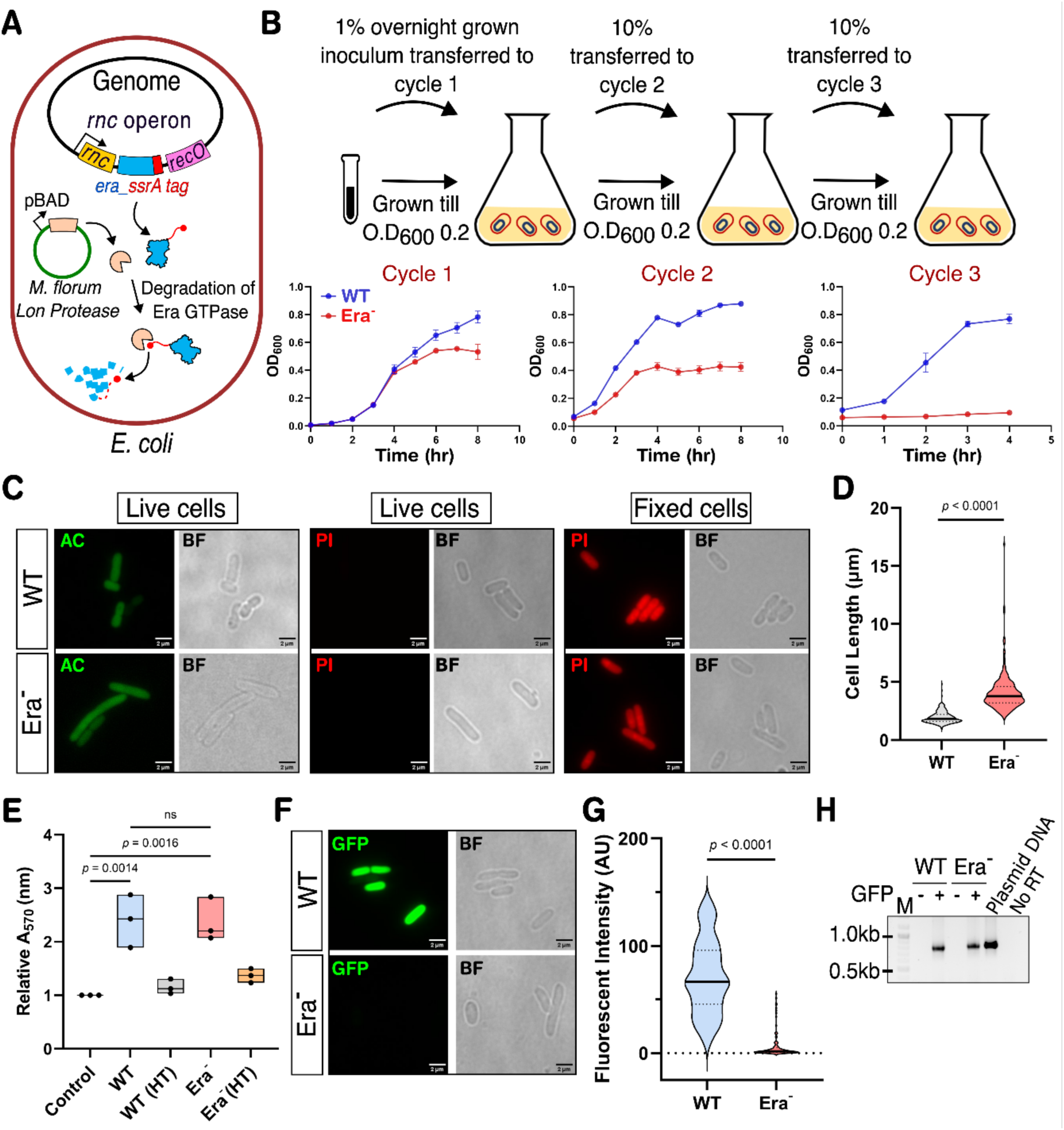
Depletion of Era leads to severe growth-associated defects in E. coli. **(A)** Schematic representation of Era depletion using *M. florum* SsrA tag and Lon protease degradation system. **(B)** Gradual depletion of Era over three consecutive growth cycles leads to growth arrest [n=3 biological replicates]. **(C)** Differential staining using Acridine orange (AC) in green and Propidium Iodide (PI) in red shows that Era^-^ cells are viable similar to WT. Fixed cells are used as control for dead cells as they are able to take up PI. **(D)** Era^-^ cells have an elongated phenotype, suggesting that they are arrested during cell division in comparison to WT [Statistical analysis: Unpaired t-test, n=3 biological replicates]. **(E)** MTT assay showed that Era^-^ cells are metabolically active similar to WT, Era^-^ heat killed (HT) cells were used as control representing metabolically inactive cells. [Statistical analysis: Ordinary One-way ANOVA, n=3 biological replicates]. **(F)** *in vivo* translation assay using GFP based reporter system showed that Era^-^ cells are not competent for protein synthesis compared to WT. **(G)** The fluorescent intensity from each bacterial cell (N = 100) from (F) has been estimated relative to cell area and represented by a violin plot [Statistical analysis: Unpaired t-test, n=3 biological replicates]. **(H)** RT- PCR showed that Era^-^ cells synthesise mEGFP mRNA upon induction. As a control, plasmid DNA was used as a template for PCR.

### Absence of Era leads to translational arrest *in vivo*

The cell growth arrest and dormancy in Era^-^ cells prompted us to probe the state of translation. We performed an *in vivo* translation assay in Era^-^ cells by monitoring the production of mEGFP using live-cell fluorescence microscopy. mEGFP fluorescence was drastically reduced in Era^-^ cells when compared to WT (Fig. 1F-G). Intriguingly, analysis of the mEGFP transcript with Reverse transcription PCR (RT-PCR), showed its presence in both the WT and Era^-^ cells (Fig. 1H) indicating that the RNA is synthesised but the translational machinery is compromised in the absence of Era, explaining the observed growth arrest (Fig. 1B). The impaired translation can be attributed to defects in the translational apparatus or downregulation of the translation process. Therefore, we assessed the state of ribosomes in Era^-^cells by polysome profiling using sucrose density gradient sedimentation. Overall, we didn’t observe major alteration of ribosome profile between WT and Era^-^ cells (Fig. 2A).

**Figure 2.**
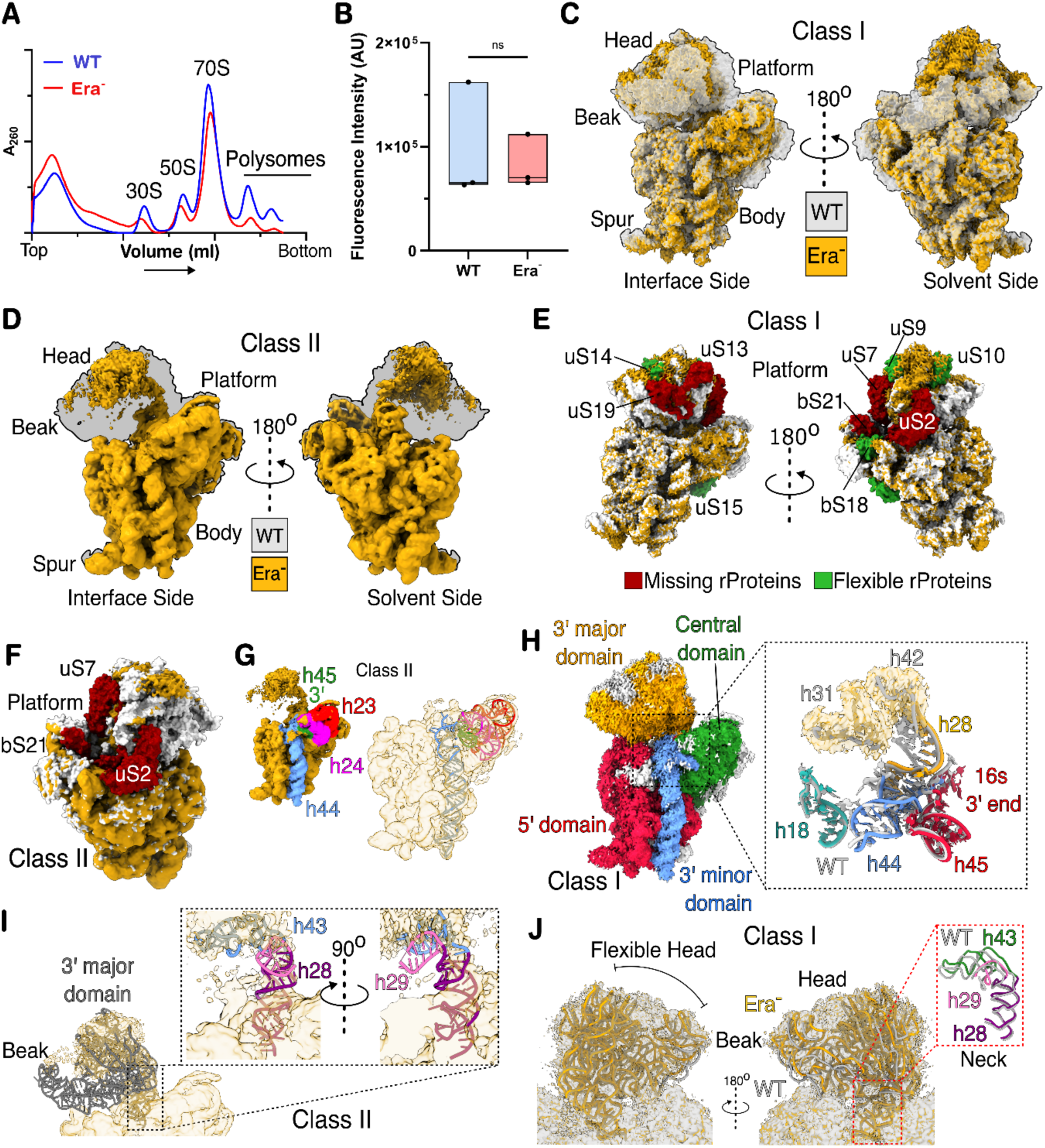
Absence of Era leads to 30S maturation defects. **(A)** Polysome profiling from Era^-^ in comparison to WT cells, showed no major alteration in the profile of monosomes or free subunits. **(B)** *in vitro* translation assay using a GFP-based reporter system showed that ribosomal subunits isolated from Era^-^ cells are functionally active, as they are able to synthesise GFP at levels similar to WT ribosomal subunits. [Statistical analysis: Unpaired t-test, n=3 biological replicates]. **(C & D)** Cryo-EM maps of 30S obtained from Era^-^ cells (depicted in yellow surface representation) showed the presence of two distinct populations; the atomic model (PDB ID: 7K00) for the WT 30S (depicted in grey surface representation) was superimposed onto the Cryo-EM density map of Class I (C) and Class II (D) to show the difference in density. **(E)** The difference between the Class I map and the WT model is highlighted by marking the missing density (red) of rProteins in the WT model, and similarly for rProteins with flexible density (green). **(F)** Similarly, in Class II, the missing (red) rProteins of the platform region are also highlighted. **(G)** The body region of Class II (surface in transparent yellow) fitted with WT model, shows helices [h23 (red), h24 (pink), h44 (blue), and h45 (green)] associated with the decoding centre are correctly folded into the mature conformation. **(H)** Representation of the different domains of 16S rRNA in the Class I map, highlighting the 5’ (red), central (green), 3’ major (yellow), and 3’ minor (blue) domains. Major rRNA helices [h18 (cyan), h28 (yellow), h44 (blue), h45 (red)] associated with the decoding centre (box) are represented in ribbon with WT model (grey) superimposed onto Era^-^ model. **(I)** 3’ domain of 16S rRNA WT model (ribbon in grey) superimposed onto the Class II map (surface in transparent yellow) shows the absence of head, indicated highly disordered conformation. **(J)** Docked 3’ domain of 16S rRNA WT model (ribbon in grey) onto the Class I map (surface in transparent yellow) shows flexible and premature conformation of the head with the helices associated with neck (inset) highlighted in ribbon in respective colours.

To further validate the functionality of ribosomes from Era^-^ cells, we isolated the ribosomes and tested their ability to synthesise proteins using an *in vitro* translation assay. For this, purified ribosomes (30S and 50S) from Era^-^ cells were added to a PURE-EXPRESS translation system and assayed for their ability to synthesise mEGFP. In contrast to the *in vivo* mEGFP reporter assay, significant fluorescence was observed with ribosomes purified from Era^-^ cells, suggesting that these ribosomes are competent for protein synthesis *in vitro* (Fig. 2B). However, the downregulation of translation *in vivo* in Era^-^ cells alludes to the possibility that there is an additional role for Era in translational regulation.

### Era depletion leads to 30S Assembly defects

The above observation of reduced translation *in vivo* but active translation *in vitro* with purified ribosomes from Era^-^ cells prompted us to analyse the structure of ribosomes. To realise this, we purified 30S subunits from Era^-^ cells and reconstructed the 3D structures using single-particle Cryo-EM. Two dominant populations of the 30S subunit, hereafter referred to as Class I and Class II, with nominal resolutions of 3.1 Å and 3.0 Å, respectively, were obtained after classification (Fig. 2C-D & S1). The density for several rProteins of the head and platform region of Class I was found to be absent (uS2, uS7, uS13, uS19 and bS21)or fragmented (uS9, uS10, uS14, uS15 and bS18), indicating low occupancy (Fig. 2E). Similar observations were also made for the Class II 30S particles, where key proteins of platform region such as uS2, uS7 and bS21 were absent (Fig. 2F). In contrast, both Class I and Class II maps showed well-resolved density for much of the rRNA helices associated with the 5’ central, and 3’ minor domains, like WT (Fig. 2G-H). However, the 3’ major domain, which comprises of the head region of the 30S, was not well resolved in both the classes with Class II being the least well defined (Fig. 2C-D, 2I-J, S1D-E). Our analysis for Class I showed that this variability in conformation of the helices associated with head is limited to the beak and lower part of the 3’ major domain (h31-34, h41-42), whereas the neck (h28-29, h43) connecting the body with the head does not show variable conformation (Fig. 2J). The 3’ end of the 16S, comprising of h45, is in a folded conformation like WT (Fig. 2H).

### 30S subunit in 70S shows heterogeneity in the absence of Era

Since we observed a prominent peak for 70S monosomes in our ribosome profile, we also performed Cryo-EM analysis of the 70S ribosomes. We found a high degree of heterogeneity in the 70S population (Fig. S2-S3 & S4). Extensive 3D classification of 70S shows that few of the 3D classes contain tRNA, despite harbouring assembly defects (Fig. 3A-B & S4B). We observed that one of the 70S subclass (Fig. 3C) associated with Class 2 has low occupancy of 30S rProteins associated with the platform (uS2 and bS21) and the position of 30S head shows differences in comparison to WT (Fig. 3D & S4B). Similarly, Class 3 also showed low occupancy for 30S rProteins associated with the platform (uS2 and bS21) but showed no presence of tRNA (Fig. 3E & S4B). In contrast to 30S, the 50S in all the 70S classes shows no maturation defects (Fig. 3 & S3-S5). We analysed the key bridge points between 30S and 50S that allow their association to form 70S. We observed that, except for minor conformational variations in B1 and B2a/d due to the flexibility of the 30S head, all other bridge points are intact (Fig. S5F-G). Thus, it is observed that in the absence of Era, the maturation of 30S head is stalled, and despite this assembly defect, the 30S is able to associate with the mature 50S to form the 70S monosome, suggesting that the maturation defect of the 30S is tolerated to form 70S. It is worth noting that the presence of tRNA (and mRNA in one class) further suggests that these 70S are likely to be competent for translation (Fig. 3A-C).

**Figure 3.**
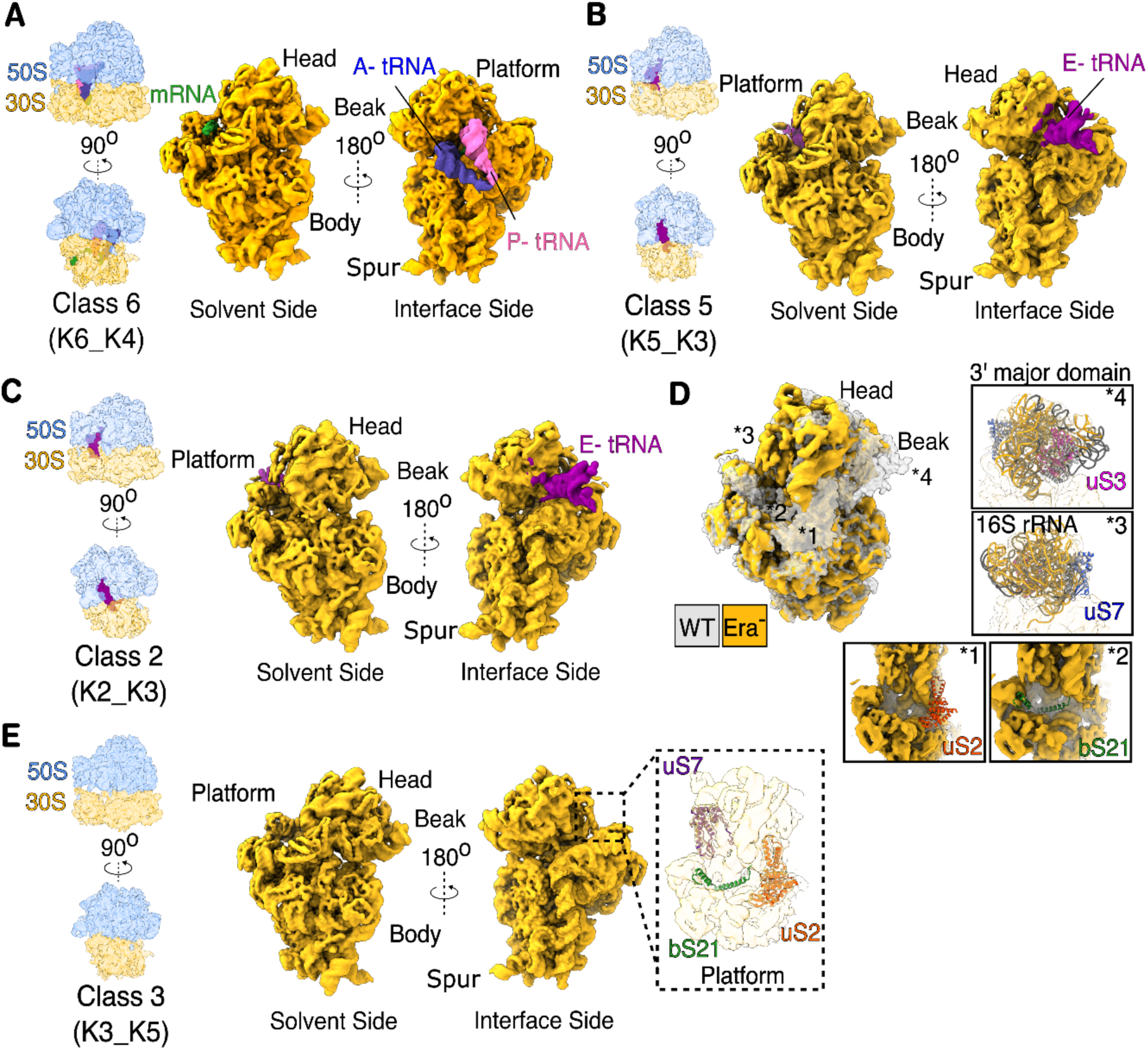
70S ribosomes from Era-depleted cells show 30S maturation defect. Cryo-EM maps of 70S ribosome from Era^-^ cells show the presence of multiple classes of 70S. **(A)** Class 6 (K6_K4) is fully mature with P- (indigo) and A-site tRNA (pink). **(B)** Class 5 (K5_K3) is near-mature, with the E-site tRNA (purple). (**C**) Class 2 (K2_K3) shows the presence of E-site tRNA (purple) but with maturation defects. (**D**) The WT 30S model (depicted in grey surface representation) superimposed onto Class 2 (K2_K3) map (depicted in yellow surface representation) shows the absence of density for uS2 (red ribbon in box 1) and bS21 (green ribbon in box 2). The rRNA helices in the head of Class 2 (K2_K3) shows an altered conformation (yellow ribbon) compared to the mature 30S (grey ribbon), based on rigid body docking (box 3 & 4). The position of uS3 (magenta ribbon in box 4) and uS7 (blue ribbon in box 3) in the head is shown and density for these proteins was observed. **(E)** Class 3 (K3-K5) contains no tRNA but shows maturation defects in the platform regions of 30S where the density for bS21 (green ribbon in insert) is found to be absent and that of uS2 (red ribbon in insert) and uS7 (purple ribbon in insert) are present.

### Era counters stringent response to promote translation and cell growth

The phenotypes observed in Era-depleted cells resemble the state of quiescence that is promoted by the stringent response in bacteria when subjected to environmental stress, such as nutrient limitation and antibiotics. Intrigued by this, we hypothesised that the absence of Era may lead to the activation of the stringent response, and that Era may be necessary to counteract the negative regulation of the stringent response on ribosome biogenesis and translation. In *E. coli*, the stringent response pathway is mediated by the effector molecule called alarmone ((p)ppGpp), synthesised primarily by RelA and counter-regulated through hydrolysis by SpoT (8, 9). Since RelA is the primary driver of the stringent response-mediated dormancy, we investigated whether the two factors, RelA and Era, are genetically linked. We created a null mutant of *relA* (Δ*relA*) using the λ-red recombineering system and depleted Era against the Δ*relA* background using *M. florum* Lon protease.

In contrast to Era^-^ cells, the growth of RelA^-^ Era^-^ cells was gradually restored to WT in subsequent cycles (Fig. 4A). This suggests that the genes coding for Era and RelA are genetically linked, and the essentiality associated with Era is contingent on the presence of RelA. This further prompted us to investigate whether the translational arrest observed *in vivo* can also be rescued in RelA^-^ Era^-^ cells. We employed β-galactosidase assay to monitor translation in RelA^-^ Era^-^ cells. This showed that RelA^-^ Era^-^ cells could synthesise β-galactosidase at a rate similar to WT, whereas Era^-^ cells showed drastically reduced rate of β-galactosidase activity, suggesting that there is a downregulation of translation in Era^-^ cells, which correlates well with impaired growth (Fig. 4B).

**Figure 4.**
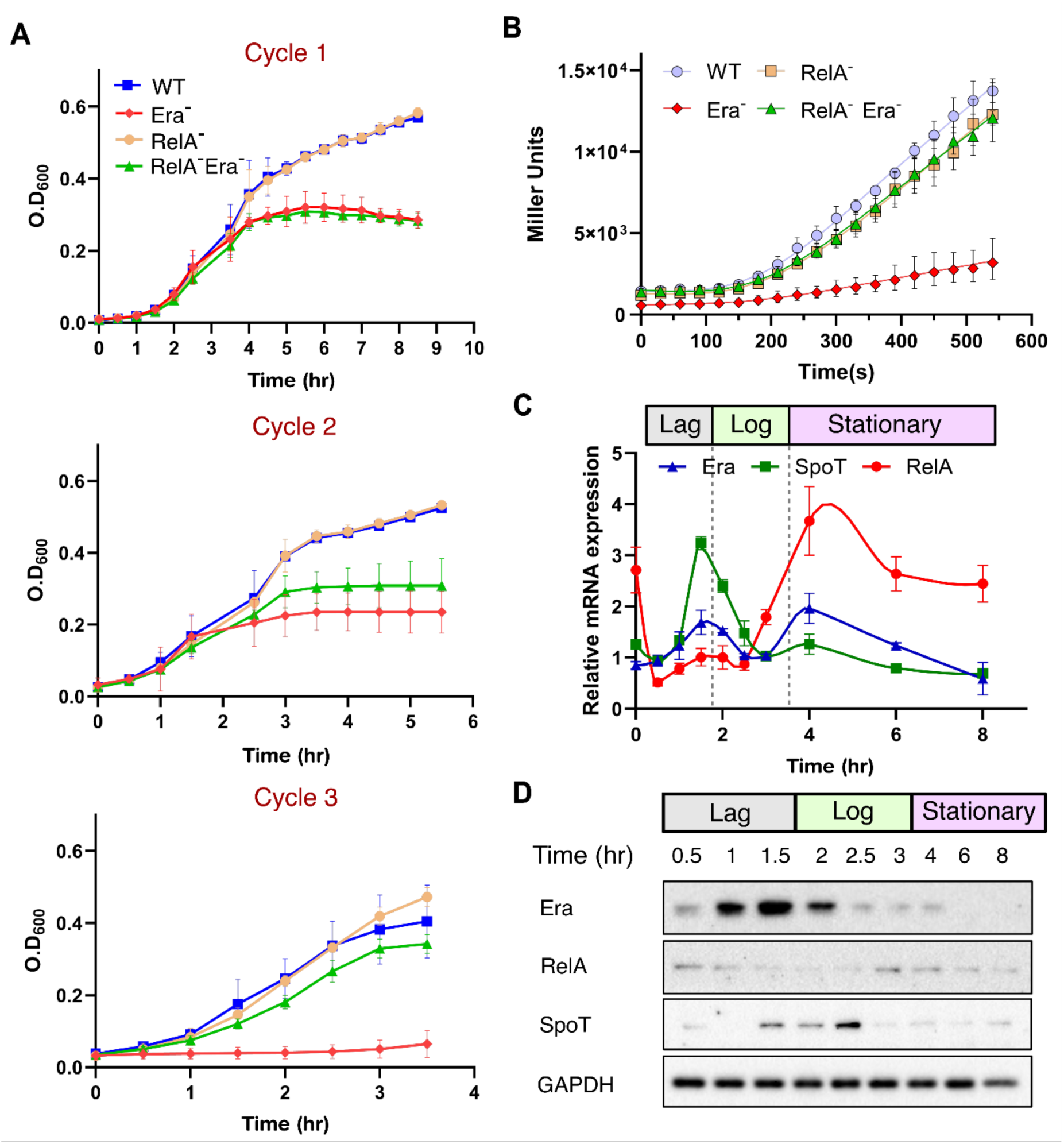
Era counteracts the effects of RelA. **(A)** Growth profile shows growth-associated defects in Era^-^ can be rescued by Era^-^ Δ*relA* [n=3 biological replicates]. **(B)** *in vivo* translation, as measured by a β-galactosidase-based reporter assay, showed that translation recovers in cells lacking both RelA and Era [n=3 biological replicates]. **(C)** mRNA expression profiles of the respective genes *era*, *relA*, and *spoT* at different time points during growth, quantified by qPCR [n=3 biological replicates]. **(D)** Representative protein production profile of Era, RelA, and SpoT at different time points during growth were detected using Western blot.

To further understand the association between Era and the factors of the stringent response, we investigated the expression profile of genes coding for Era, RelA, and SpoT across different phases of bacterial cell growth at both the mRNA and protein levels using quantitative PCR (qPCR) and Western blotting, respectively (Fig. 4C-D). We observed that production of Era and RelA is mutually exclusive in the stationary phase, where synthesis of RelA was upregulated but not Era. Whereas, in the log phase, both Era and RelA are downregulated. Interestingly, in the lag phase, we found that the production of Era and RelA shows anti-correlation, wherein the production of Era rises, but that of RelA declines (Fig. 4C-D). We observed that SpoT, which is a (p)ppGpp hydrolase, is only produced in the log phase, which led us to hypothesise that in the absence of SpoT, Era is necessary to counteract RelA in the lag phase, and allow the cells to transition towards exponential growth. In line with the protein profile, the mRNA expression profile of Era, RelA and SpoT generally follows similar trend; however, we observed a second peak for Era at the onset of stationary phase, which declines thereafter, correlating with the protein level (Fig. 4C-D).

### Era downregulates translation

Encouraged by the observation that Era counteracts stringent response to promote translation and cell growth, we investigated the underlying mechanism. As a first step, we probed the state of ribosomes in lag phase cells, and this showed reduced 70S monosomes with a concomitant increase in free subunits (Fig. 5A). We observed that Era tends to co-migrate with free 30S (Fig. 5A). Since (p)ppGpp is a known modulator of cellular GTPases involved in ribosome biogenesis and translation (10–12), we tested the hypothesis that Era can interact with alarmones directly, sequestering them to preclude downregulation of translation. Using the *in vitro* translation system with mEGFP as the readout, we observed that the addition of (p)ppGpp reduced the total GFP output (Fig. S6), an effect that we attempted to rescue by adding Era (Fig. S6), in accordance with our proposed hypothesis. However, we found that Era tends to downregulate translation on its own (Fig. 5B). This surprising observation prompted us to ask how Era inhibits translation. Based on 30S binding both *in vivo* (Fig. 5A) and *in vitro* (Fig. 5C), we hypothesised that Era may interfere with 70S formation. This alludes to a scenario where Era might maintain subunits in a free state by either dissociating 70S or preventing the association of 30S and 50S to form 70S. However, unlike ObgE and HflX GTPases (39, 40), that are known to dissociate 70S, we observed no such activity for Era (Fig. 5D-E), despite its tendency to interact with 30S (K_d_ = 0.6929 µM), as probed by the co-sedimentation assay (Fig. 5C).

**Figure 5.**
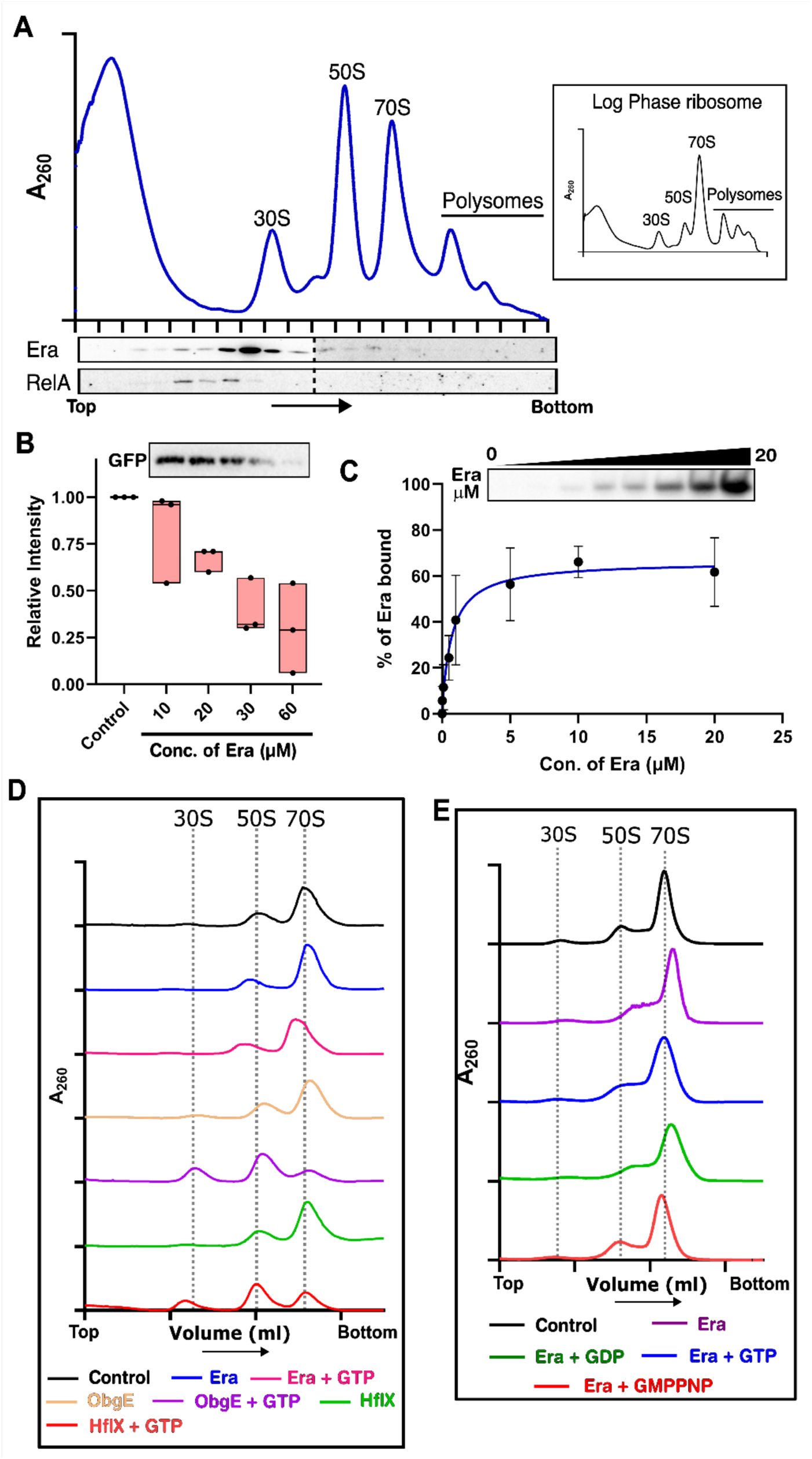
Effect of Era on ribosome and translation. (**A**) Polysome profiling from the lag phase cells shows a higher amount of free subunits in comparison to 70S monosomes. Representative ribosome profile from log phase cells are depicted in inset for comparison. Representative western blot analysis of ribosome fractions showed Era tends to co-migrate with the 30S. **(B)** *An in* vitro translation assay using a GFP-based reporter system showed that the presence of Era inhibits translation. A representative blot corresponding to GFP signal is depicted. Western blot and densitometric analysis were used to quantify the GFP [n=3 biological replicates]. **(C)** *in vitro* co-sedimentation assay using purified 30S and Era followed by western blot and densitometric analysis showed that Era binds to 30S with a K_d_ of 0.6929 µM [n=3 biological replicates]. A representative blot of the pellet fraction corresponding to the signal intensity of 30S-bound Era is shown. See methods for details. **(D)** The sucrose density gradient profile showed that Era is unable to dissociate the mature 70S ribosome, unlike the GTPases ObgE and HflX. **(E)** The sucrose density gradient profile monitoring the association of subunits suggests that Era does not prevent the association of 30S and 50S to form 70S ribosome.

### Era rescues translating ribosomes from stringent response

In the absence of any known dissociative or associative activity, we proceeded to investigate other modes by which Era might inhibit translation. In response to cellular stress, RelA has been shown to bind to 70S in the presence of deacylated tRNA at the A site. This stimulates the synthetase domain, producing (p)ppGpp, which initiates the stringent response (41–45). Encouraged by the genetic interaction between Era and RelA, we tested that whether Era binds directly to RelA. We used protein-protein immunoprecipitation to track whether Era can pull-down RelA *in vitro* when used as a bait. We indeed observed a positive interaction as Era was able to pull-down RelA, which was analysed by Western blotting using an anti-RelA antibody (Fig. 6A). The binding isothermin the absence (K_d_ = 0.863 µM, n_H_ = 2.709) and presence (K_d_ = 0.8966 µM, n_H_ = 4.138) of GMPPNP showed comparable affinity and cooperative binding indicating the possibility of more than one binding site (Fig. S7B). To assess the role of G (1-185 aa) and KH (186-301 aa) domains in RelA binding, we cloned and purified these domains separately and tested their interaction with RelA. We found that G-domain (K_d_ = 0.523 µM, n_H_ = 2.770) and KH-domain (K_d_ =0.441 µM, n_H_ = 6.248) can independently interact with RelA, respectively, with comparable binding affinity (Fig. S7A). Thus, in line with the genetic interaction, Era and RelA also exhibit physical interaction, which raises a question on their combined role in protein synthesis. To clarify this, we added RelA and (p)ppGpp to the *in vitro* translation reaction and noticed a reduction in protein synthesis. In contrast to this, when Era was added to this reaction, we observed that protein synthesis is restored to a level comparable to that of the control (Fig. 6B). This suggests that although RelA and Era individually suppress protein synthesis through distinct mechanisms, their combined effect nullifies their individual inhibitory activities.

**Figure 6.**
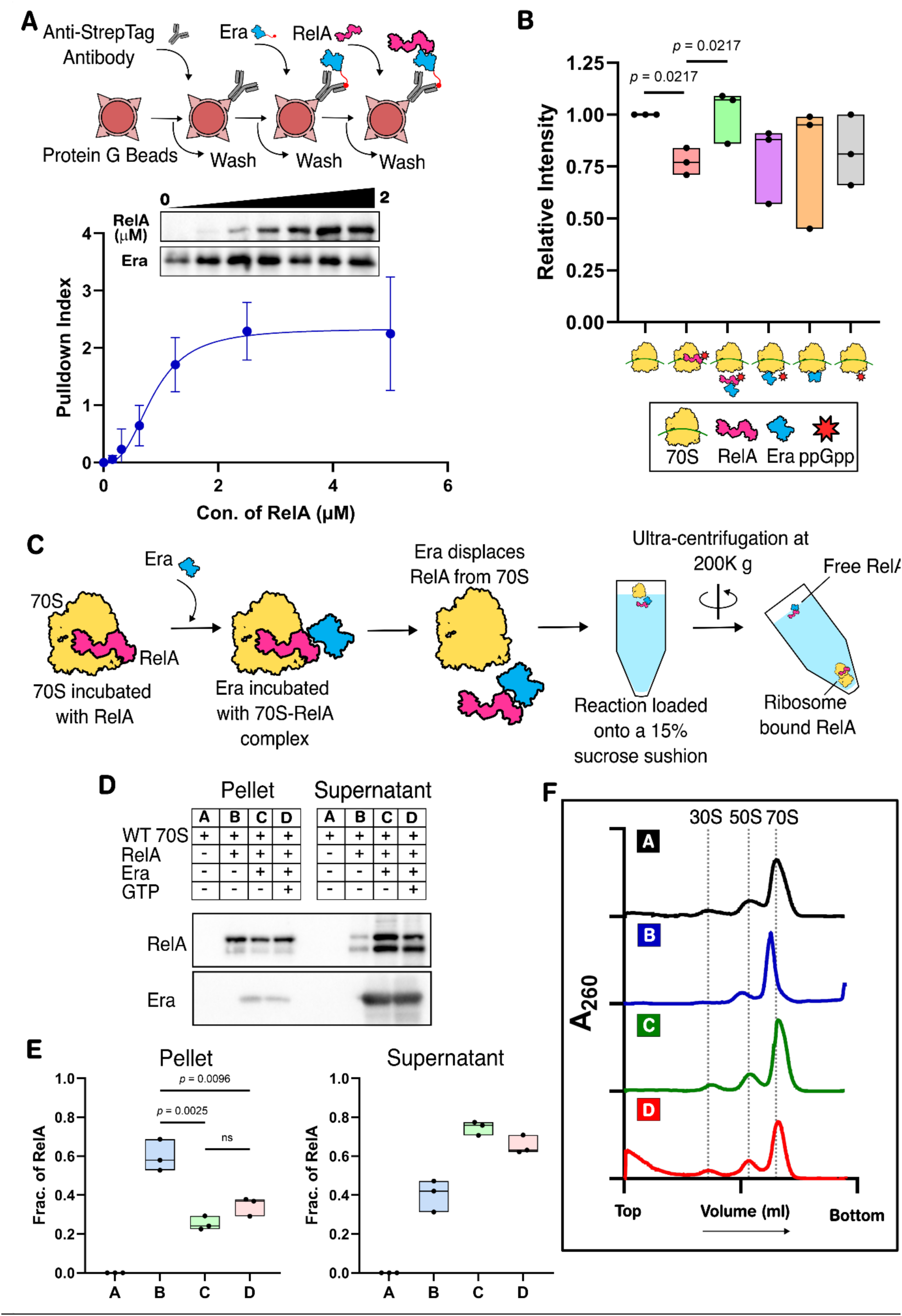
Era rescues ribosomes from RelA-mediated inhibition. **(A)** Interaction between Era and RelA was probed using a protein-protein pull-down assay as described in the representative schema. The binding was analysed using Western blot and densitometry to estimate the dissociation constant (K_d_ = 0.863 µM, n_H_ = 2.709, R^2^ = 0.844, n=3 biological replicates). **(B)** *in vitro* translation assay using a GFP-based reporter system showed inhibition of translation in the presence of RelA, which was rescued upon addition of Era [Statistical analysis: Ordinary one-way ANOVA, n=3 biological replicates]. **(C)** Representative schema showing the experimental assay to assess the release of 70S-bound RelA by Era. **(D)** Western blot shows that Era releases RelA bound to 70S ribosomes. **(E)** Western blot data (D) were analysed by densitometry to determine the amount of RelA in the pellet and supernatant fractions [Statistical analysis: Unpaired t-test, n=3 biological replicates]. **(F)** Co-sedimentation assay shows no changes to the ribosome profile on addition of RelA and subsequent release of RelA by Era.

We envisaged two possibilities to explain how the combination of Era and RelA can restore protein synthesis. i) Era sequesters RelA to prevent it from binding to 70S, and ii) Era may bind to 70S-RelA complex and release RelA from it. The physical interaction between Era and RelA alluded to the first scenario of RelA sequestration (Fig. 6A), and we tested the second possibility using ribosome pelleting assay. Purified 70S ribosomes were pre-incubated with RelA at an equimolar ratio to allow RelA binding to the 70S ribosomes. This binding was then challenged with a fivefold excess of Era. Following incubation, the reactions were ultracentrifuged to pellet the ribosomes. The free (supernatant) and ribosome-bound RelA (pellet) were quantified by Western blotting. The amount of RelA pelleting with ribosomes decreases by about two-thirds once challenged by Era (Fig. 6D-E). This suggests that Era promotes dissociation of RelA from 70S, and this is independent of GTP binding, as K21A mutant (46) of Era that abrogates GTP binding also releases RelA (Fig. 6D-E & S8A). Similar observations were also made when RelA displacement activity was tested with individual domains of Era. Although both the G-domain and KH-domain can independently release RelA, the G-domain exhibited significantly higher displacement activity than the KH-domain. (Fig. S8B-C). To assess whether the release of RelA from 70S by Era leads to a change in the composition of 70S,the reaction mixtures were subjected to polysome profiling. This showed that RelA displacement from 70S does not lead to dissociation of subunits (Fig. 6F). These assays suggest that Era displaces 70S-bound RelA without splitting the 70S and thereby promotes protein synthesis.

## Discussion

Adaptation to a changing environment is a fundamental attribute for the survival of bacterial cells. In response to favourable environmental cues, cells transition into an active growth phase, whereas an environmental stress promotes entry into a dormant state. While the mechanisms governing the transition from growth to dormancy are reasonably well understood, the processes underlying exit from dormancy remain poorly characterised. The dormant-like state that ensues in the absence of Era GTPase allowed us to investigate the regulatory mechanism for cellular resuscitation (Fig. 1). We find that the Era-depleted cells are viable and metabolically active (Fig. 1C-E). These characteristics are similar to the cells that enter dormancy upon induction of stress. Further investigations revealed that the protein synthesis is downregulated in Era-depleted cells (Fig. 1F-G). Since Era is involved in 30S maturation (16–18), we reasoned that any assembly defect in Era-depleted cells might curtail 30S maturation, thereby impacting protein synthesis. While the assembly defect was not apparent from the polysome profiling (Fig. 2A), the Cryo-EM analysis of 30S from Era-depleted cells revealed a highly flexible head and varying occupancy for several rProteins (uS2, uS7 and bS21) in the platform region (Fig. 2C-J). This suggested that the arrest of protein synthesis in Era-depleted cells could be due to 30S assembly defects. However, the decoding centre in 30S is fully formed and shows features like the WT (Fig. 2H). We also note that, unlike the 3’ major domain of 16S rRNA (head region), the 5’, central, and 3’ minor domains are folded, suggesting that these domains can fold to their mature state independent of the 3’ major domains (Fig. 2H). This concurs with the earlier report that suggests that whenever folding of a particular rRNA domain is stalled, the folding of other rRNA domains can be rerouted through multiple parallel pathways, leading to co-maturation (18). Surprisingly, the 30S and 50S ribosomes purified from Era-depleted cells were competent for protein synthesis *in vitro*, suggesting that these maturation defects are tolerated (Fig. 2B). In line with this, the Cryo-EM analysis of 70S ribosomes from Era-depleted cells showed that the 30S and 50S association remains intact (Fig. 3 & S5F-G). Though the 70S data set showed high heterogeneity (Fig. 3, S2 & S3), the components of the 50S were fully assembled in all the classes. Whereas in 30S, some classes showed assembly defects, wherein some rProteins are absent (uS2 and bS21) and exhibit variable occupancy (uS7 and uS19). Despite this, most of the bridge points (except B1 and B2a/d) between the subunits were intact (Fig. S5G), and in some 70S subclasses with 30S assembly defects, tRNAs were also observed (Fig. S4B), suggesting that the 70S retain translational competence despite the 30S assembly defects in the absence of Era (Fig. 3). This further suggests that the selective downregulation of translation *in vivo* in Era-depleted cells occurs through an alternative process.

RA-GTPases have been shown to interact with factors of the stringent response pathway (10–12). *E. coli* has two RSH family of enzymes involved in modulating the stringent response pathway. The long monofunctional RelA is primarily involved in the synthesis of (p)ppGpp, which serves as an alarmone to upregulate the stringent response during nutrient starvation, whereas the bifunctional SpoT shows a higher hydrolase activity towards alarmone than the synthetase activity (8, 9). Era GTPases have been shown to bind to (p)ppGpp with high affinity, thereby impacting the activity of Era in 30S maturation (12, 47). In *Staphylococcus aureus*, Era interacts with Rel protein, as part of a complex along with CshA, during ribosome maturation (48). Similarly, a conserved and essential RA-GTPase, ObgE, which is involved in 50S assembly, has been shown to interact with SpoT to modulate its activity (49–51). This hints at the link between ribosome maturation and stringent response pathways. In line with this, we observed that while Era-depletion leads to a dormant-like phenotype, removal of both Era and RelA effectively rescued defects in both growth and protein synthesis (Fig. 4A-B). This suggests that *era* and *relA* are genetically linked, and the essentiality of Era is contingent on the presence of RelA in *E. coli*. The fact that the essential function of Era can become dispensable upon the loss of RelA strongly suggests that Era counteracts the negative regulation of the stringent response factors on ribosome biogenesis and translation. However, this context-dependent essentiality of Era appears to change over the course of evolution, as Era is essential in some bacterial species like *E. coli* but not in others such as *S. aureus* and *Mycobacterium tuberculosis* (14, 19, 48, 52). In agreement with the genetic link between *era* and *relA*, synthesis of Era rises at the lag phase and early log phase, coinciding with the traces of RelA in lag phase (Fig. 4C-D). Incidentally, the synthesis of Era is timed just before the onset of log phase to support the rapid ribosome synthesis activity in the exponential growth phase, where RelA synthesis is not observed (Fig. 4C-D). This lends further support that Era counteracts the effect of RelA to drive ribosome synthesis and translation that are essential to launch the exponential growth. How does Era counteract the downregulation of translation by RelA? We find that Era could rescue translation from RelA-mediated inhibition *in vitro* (Fig. 6B), which is counterintuitive, as Era itself downregulates protein synthesis (Fig. 5B). Previous studies proposed that Era acts as a regulatory checkpoint blocking premature translation initiation by the immature 30S subunit (16–18). The proposed mechanism includes occlusion of the anti-Shine-Dalgarno (SD) sequence, dissociative activity on 70S ribosomes, or blocking the 50S subunit from associating with the 30S subunit (17, 18). Although we could not observe the dissociative and anti-associative activity of Era in our experiments (Fig. 5D-E), we did observe active binding of Era to 30S, both *in vitro* and *in vivo* (Fig. 5A & 5C). During nutrient limitation, RelA binding to 70S ribosome gets stabilised in the presence of deacetylated tRNA at the A-site (41, 43, 44). This activates RelA to synthesise (p)ppGpp, which then promotes stringent response leading to dormancy. This is typically seen in the stationary phase of bacterial growth, where we have also observed the synthesis of RelA, but not Era (Fig. 4C-D). Thus, it is evident that when the cells are relieved of stress (under nutrient rich condition), the 70S bound RelA needs to be released to restart protein synthesis to support growth. In this context, the role of Era, whose synthesis surges just before the onset of log phase, is to dislodge RelA from the 70S ribosome, thereby relieving the 70S from the RelA-mediated inhibition. Indeed, we observed that Era was able to release RelA from 70S both in the presence and absence of GTP (Fig. 6C-F). This release appears to be driven by the physical interaction between Era and RelA (Fig. 6A), primarily through the G-domain of Era (Fig. S7 & S8). With these observations, we propose a probable model for Era-mediated cellular resuscitation. During stationary phase where the nutrients are deprived, the presence of RelA leads to inactivation of 70S ribosome and production of (p)ppGpp, which then promotes stringent response leading to dormancy (Fig. 4C-D). However, this repression of protein synthesis is likely to continue even when the nutrients are replenished due to the presence of RelA in the lag phase (Fig. 4C-D). At this stage, the sudden surge in the synthesis of Era may result in dislodging RelA from 70S ribosome that may allow it to resume protein synthesis. At the same time, binding of Era to RelA may preclude RelA from binding to 70S again (Fig. 4D & 6D-E). Since there is no substantial synthesis of RelA and Era in log phase, subsequent cell division may dilute out RelA and Era, and the simultaneous synthesis of SpoT may lead to hydrolysis of (p)ppGpp (Fig. 4C-D). This will promote exponential cell growth until the advent of nutrient starvation. It appears that at high concentration, Era may act as an indiscriminate release factor inhibiting protein synthesis (Fig. 5B). This explains the lack of Era during the log phase (Fig. 4D). It is tempting to speculate that Era may release a 30S-bound assembly factor from head and platform region to accelerate the assembly of rProteins. This study reveals the bifunctional role of Era, wherein it not only promotes the maturation of head and platform in the nascent 30S but also participates in translational de-repression by removing RelA from 70S (Fig, 7). Our work highlights how Era counters the activation of the stringent response and allows the cell to transition toward exponential growth thus maintaining cellular homeostasis between growth and quiescence (Fig. 7). We also anticipate similar roles for the other conserved RA-GTPases in maintaining cellular homeostasis by forming a key link between ribosome biogenesis and stringent response. This further opens the possibility of investigating the potential role of these GTPases in regulating bacterial growth and adaptation in response to changing environmental conditions.

**Figure 7.**
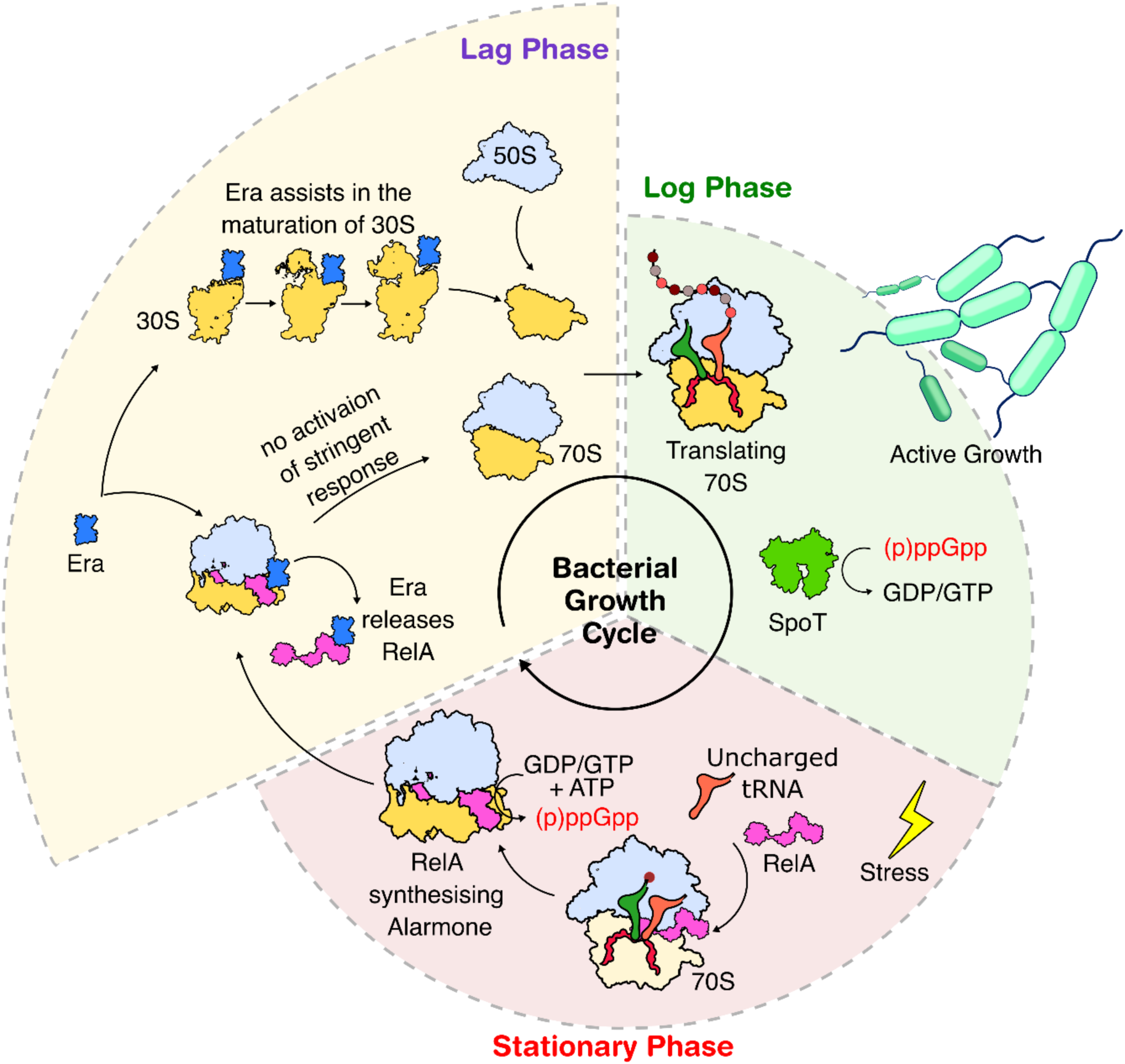
Bifunctional role of Era in cellular homeostasis. A schematic model shows the bifunctional role of Era in 30S maturation and rescue of 70S from RelA-mediated downregulation to promote growth during the log phase of bacterial growth.

## Data Availability

The 30S Class I model is deposited to PDB [29VN]. The EM maps for 30S Class I [EMD-57399], 30S Class II [EMD-80229], 70S Class 2 K2_K3 [EMD-80230], 70S Class 3 K3_K5 [EMD-80231], 70S Class 5 K5_K3 [EMD-80232] and 70S Class 6 K6_K4 [EMD-80233] are deposited to EMDB.

## Acknowledgement

Vector pET-1R (Addgene #29664), pET-13S-R (Addgene #48328) and pET His6 GFP (1GFP) (Addgene #29663) was a kind gift from Scott Gradia. Vector pLJSRSF7 (Addgene #64693) was a kind gift from Hideo Iwai. Vector pBAD33-mf-lon (Addgene #21867) was a kind gift from Robert Sauer. We thank all members of the MAB lab for their critical comments and suggestions.

## Author contribution

RP and BA conceptualised the study. RP, HM, and BA designed the experiments. RP, HM, PS, and KRV performed the experiments. RP, HM, PS, KRV, and BA analysed the data. BA acquired funding for the project. RP and BA wrote the paper with inputs from HM and KRV, while all authors reviewed at least one draft.

