## Supplementary Material for "Epistatic interaction between ribosome-associated Era GTPase and stringent response regulator RelA modulates bacterial cell growth and dormancy"

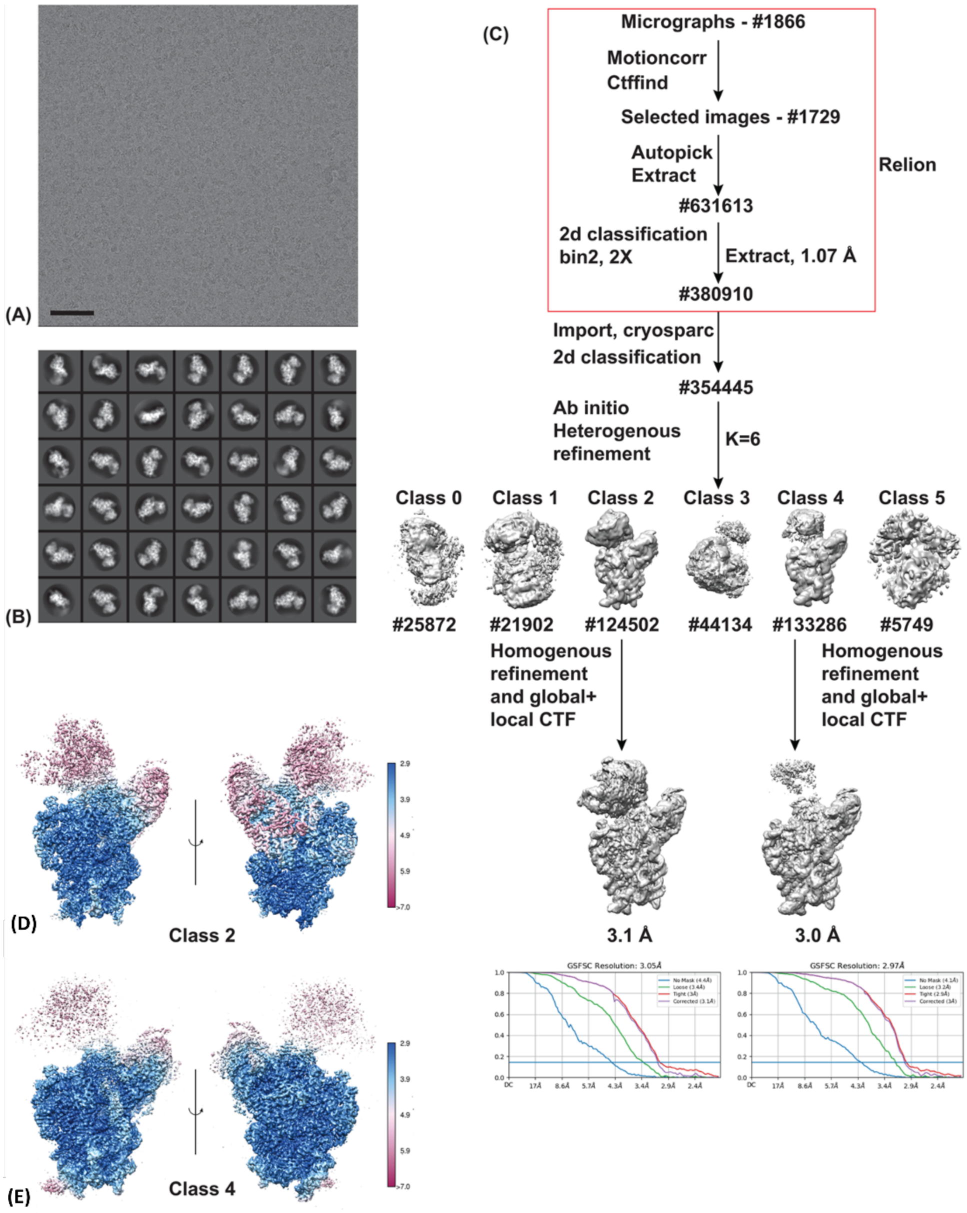

**Figure S1.** ***Cryo-EM workflow of the 30S ribosomal subunit image processing from Era-depleted (Era⁻) cells***

**(A)** A representative micrograph of 30S from Era⁻ cells shows distinct views of the molecule (scale bar – 500 Å). **(B)** Select 2D class averages of 30S are shown; box size is 160 pixels and 2.14 Å pixel size. **(C)** The Cryo-EM workflow of 30S Era⁻ data set is shown. The steps denoted in red rectangle were performed with Relion 4.0 and the rest with CryoSPARC . Two major classes with well-defined body of 30S was observed, while the head and the beak domains are less well ordered. The final unsharpened map (grey) of the two classes are shown, and the gold-standard FSC from CryoSPARC indicates resolutions of 3.1 Å and 3.0 Å, respectively, for the two classes. **(D & E)** Local resolution plots of the two classes (Class 2 and Class 4) of Era$^-$ 30S are shown. As expected, the body of the 30S is well-resolved, but the head and beak are at lower resolution. In the main text, Class 2 is denoted as Class I and Class 4 is denoted as Class II.

***Table S1: Statistics of the 30S Class I Model***

| **Model (PDB: 29VN)/ Map (EMDB: EMD-57399)** | |
|---|---|
| **Composition (#)** | |
| Chains | 11 |
| Atoms | 29927 (Hydrogens: 0) |
| Residues | Protein: 898 Nucleotide: 1065 |
| Water | 0 |
| Ligands | MG: 32 |
| **Bonds (RMSD)** | |
| Length (Å) (# > 4σ) | 0.004 (0) |
| Angles (°) (# > 4σ) | 0.754 (11) |
| **MolProbity score** | 1.95 |
| **Clash score** | 4.22 |
| **Ramachandran plot (%)** | |
| Outliers | 0.00 |
| Allowed | 7.03 |
| Favored | 92.97 |
| **Rama-Z (Ramachandran plot Z-score, RMSD)** | |
| whole (N = 882) | -1.91 (0.27) |
| helix (N = 289) | -0.34 (0.30) |
| sheet (N = 166) | -1.73 (0.40) |
| loop (N = 427) | -1.67 (0.28) |
| **Rotamer outliers (%)** | 2.45 |
| **Cβ outliers (%)** | NA |
| **Peptide plane (%)** | |
| Cis proline/general | 0.0/0.0 |
| Twisted proline/general | 0.0/0.0 |
| **CaBLAM outliers (%)** | 3.00 |

**ADP (B-factors)**

| | |
|---|---|
| Iso/Aniso (#) | 29927/0 |
| min/max/mean | |
| Protein | 31.27/138.84/78.38 |
| Nucleotide | 0.00/367.41/82.38 |
| Ligand | 5.40/83.41/46.49 |
| Water | --- |

**Occupancy**

| | |
|---|---|
| Mean | 1.00 |
| occ = 1 (%) | 100.00 |
| 0 < occ < 1 (%) | 0.00 |
| occ > 1 (%) | 0.00 |

### Data

=================================================================

**Box**

| | | |
|---|---|---|
| Lengths (Å) | | 120.91, 159.43, 195.81 |
| Angles (°) | | 90.00, 90.00, 90.00 |
| **Supplied Resolution (Å)** | | 3.1 |
| **Resolution Estimates (Å)** | **Masked** | **Unmasked** |
| d FSC (half maps; 0.143) | --- | --- |
| d 99 (full/half1/half2) | 3.3/---/--- | 3.2/---/--- |
| d model | 3.1 | 3.1 |
| d FSC model (0/0.143/0.5) | 3.0/3.0/3.5 | 3.0/3.1/3.8 |
| **Map min/max/mean** | | 2.62/6.51/0.19 |

### Model vs. Data

=================================================================

| | |
|---|---|
| CC (mask) | 0.80 |
| CC (box) | 0.76 |
| CC (peaks) | 0.68 |
| CC (volume) | 0.81 |
| Mean CC for ligands | 0.78 |

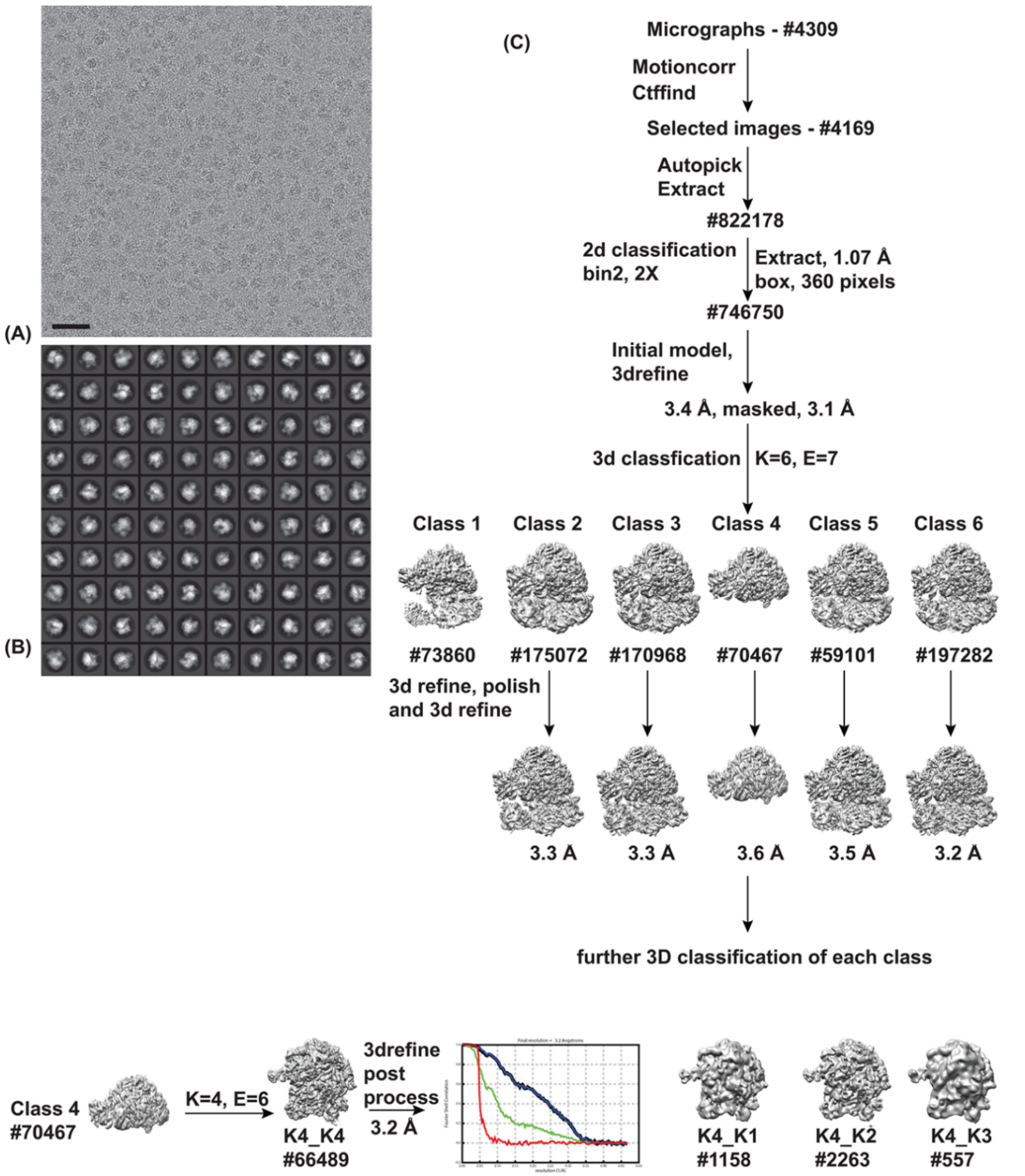

(A)
(B)
(C)
Micrographs - #4309
Motioncorr
Ctffind
Selected images - #4169
Autopick
Extract
#822178
2d classification
bin2, 2X
Extract, 1.07 Å
box, 360 pixels
#746750
Initial model,
3drefine
3.4 Å, masked, 3.1 Å
3d classfication
K=6, E=7
Class 1
Class 2
Class 3
Class 4
Class 5
Class 6
#73860
#175072
#170968
#70467
#59101
#197282
3d refine, polish
and 3d refine
3.3 Å
3.3 Å
3.6 Å
3.5 Å
3.2 Å
further 3D classification of each class
Class 4
#70467
K=4, E=6
K4_K4
#66489
3drefine
post
process
3.2 Å
K4_K1
#1158
K4_K2
#2263
K4_K3
#557

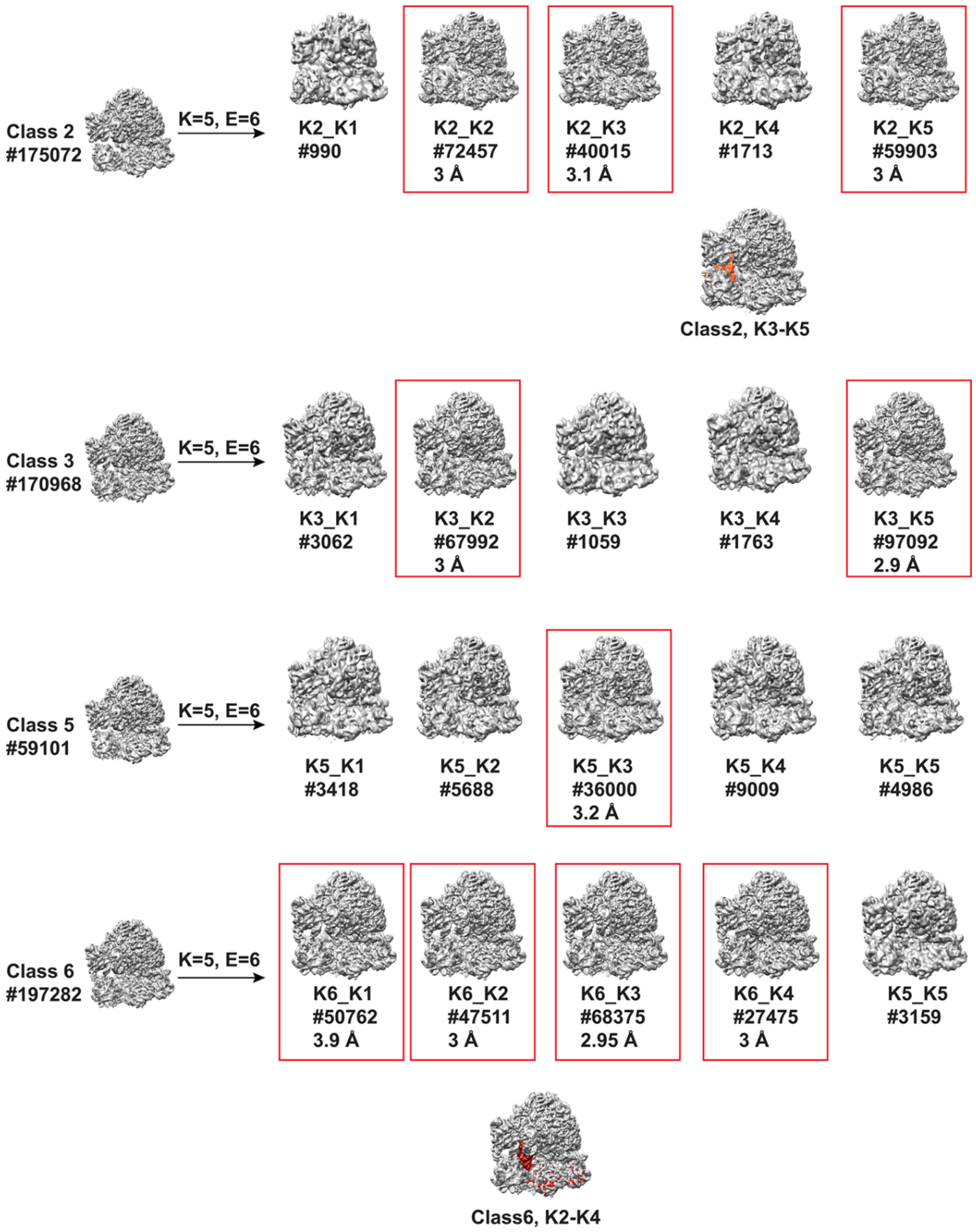

**Figure S2.** ***Cryo-EM analysis of 70S from Era⁻ cells***

(**A**) A representative micrograph of 70S from Era$^-$ cells show distinct views of the molecule (scale bar – 500 Å). (**B**) Select 2D class averages of 70S are shown; box size is 120 pixels and 3.21 Å sampling. (**C**) The Cryo-EM workflow of the 70S Era$^-$ data set is shown. All the steps were performed in RELION 4. Multiple populations of the ribosomes were identified by classification and the ones marked in red rectangle were chosen for further analysis. The presence of tRNA in few classes are shown by calculating a difference map from within the same class. For example, in Class 2, the unsharpened map from sub-class 3 and 5 were subtracted in ChimeraX and the tRNA density is shown in red. Similarly, in Class 6, the unsharpened map from sub-class 2 and 4 were subtracted in ChimeraX and the tRNA density is shown in red. The sub-class 4 has two tRNAs bound.

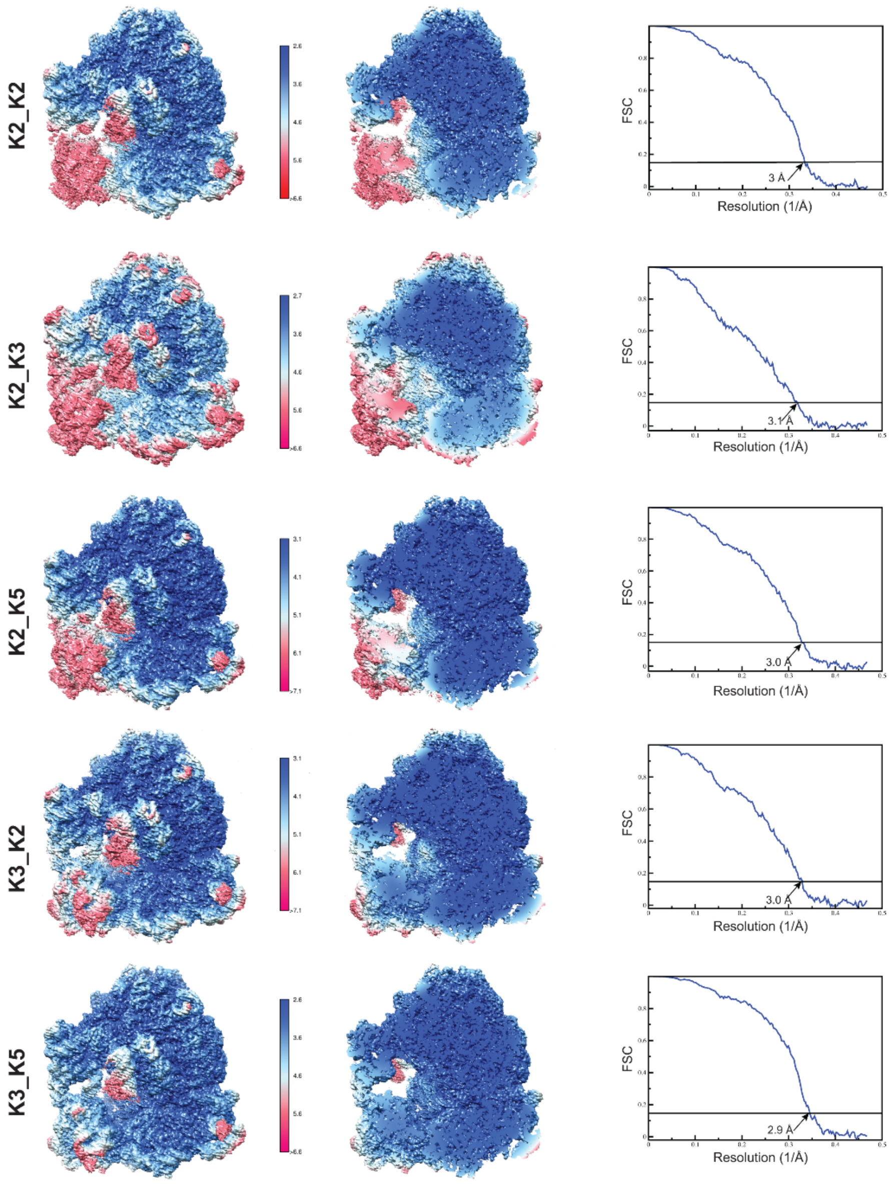

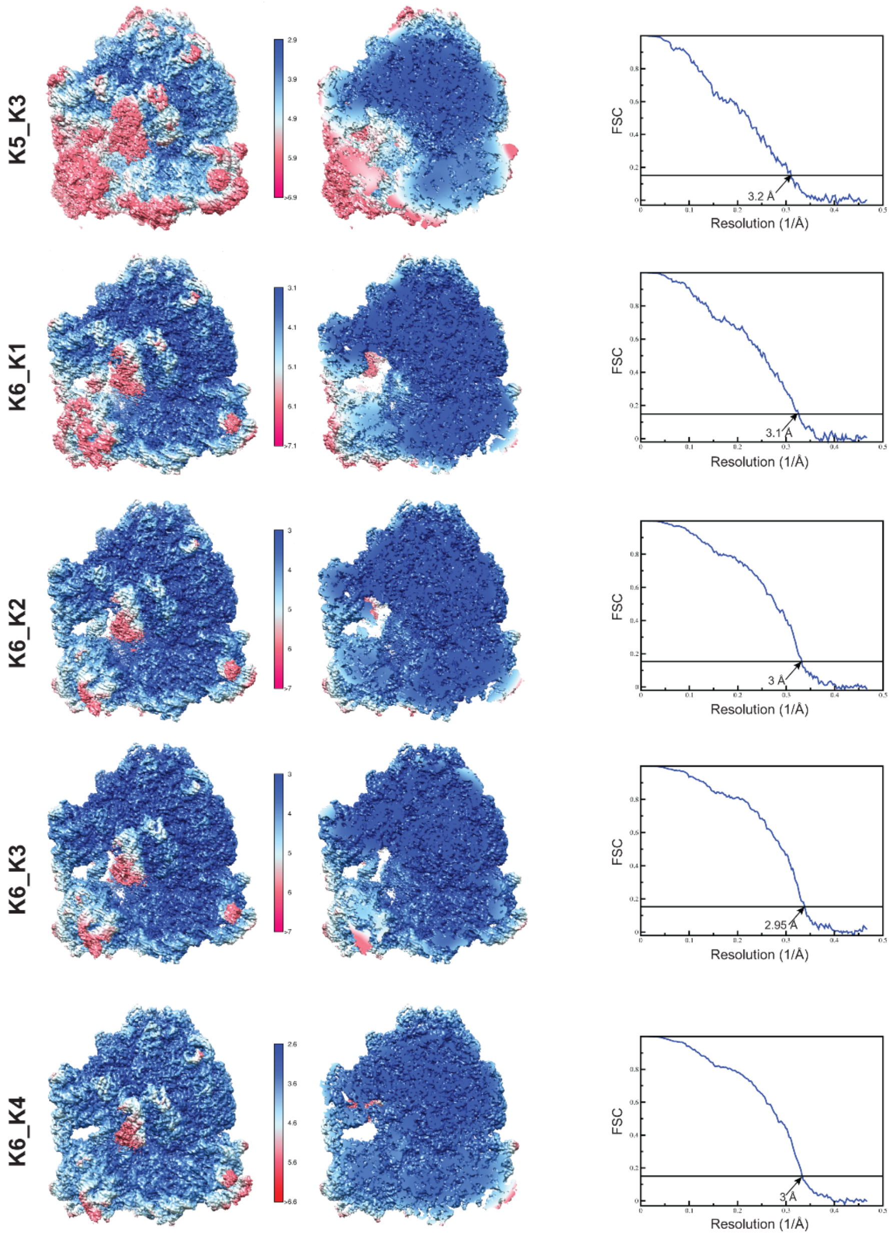

**Figure S3:** ***Local resolution plots and resolution estimate of different populations of the 70S Era⁻ ribosomes***

For each class, the local resolution plot is shown for the whole map and a slice to show the interior revealing that the large subunit and the body of the small subunit are better resolved and the head domain of the small subunit and periphery of the ribosome is less well resolved. The FSC curves calculated from two half-maps and masking from the postprocess step in RELION 4 are shown.

A

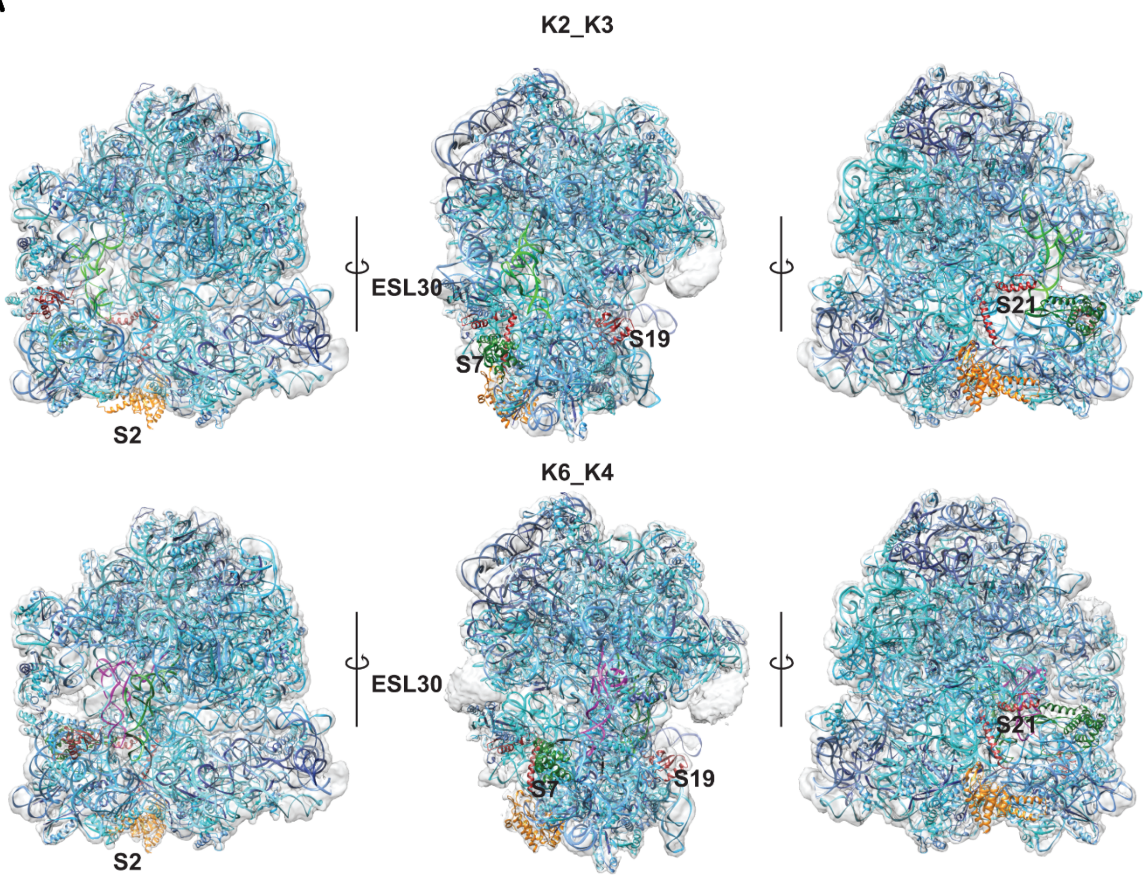

B

| Class | Missing/poor subunits | tRNA | Population (%) |
|---|---|---|---|
| K2_K2 | S2, S21, poor S7, S19 | - | 9.7 |
| K2_K3 | S2, S21, poor S7, S19 | E-site | 5.3 |
| K2_K5 | S2, S21, poor S7, S19 | - | 8.02 |
| K3_K2 | S21, poor S2 | - | 9.1 |
| K3_K5 | S21, poor S2, | - | 13 |
| K5_K3 | partial S21, poor S2,S7 | E-site | 4.8 |
| K6_K1 | S21, poor S2 | - | 6.79 |
| K6_K2 | S21, poor S2 | - | 6.36 |
| K6_K3 | S21, poor S2, S7 | - | 9.15 |
| K6_K4 | - | P+A-site | 3.67 |

**Figure S4. 3D *classes from 70S ribosomes from Era - cells***

**A)** The Cryo-EM maps of Era$^-$ 70S ribosome subclasses **K2_K3** and **K6_K4 (**in gray**)**, fitted with the wild-type model (PDB: 7K00). Subunits that are absent or display poor density in subclass **K2_K3** are highlighted in distinct colors and shown alongside the fully mature **K6_K4** for comparison. Three different views of the 70S ribosomes are shown to highlight the differences. The ESL30 of 23S rRNA in K2_K3 moves in towards the E-site tRNA, while it is unmodelled in K6_K4 due to less resolved density. The unsharpened maps after refinement are overlaid with the model colored in shades of blue.  In both classes, parts of rRNA that are not modelled can be seen as grey density with no model in them. **(B)** Table summarizing the Cryo-EM subclasses, indicating the missing or poorly resolved subunits and the corresponding percentage population of particles in each class

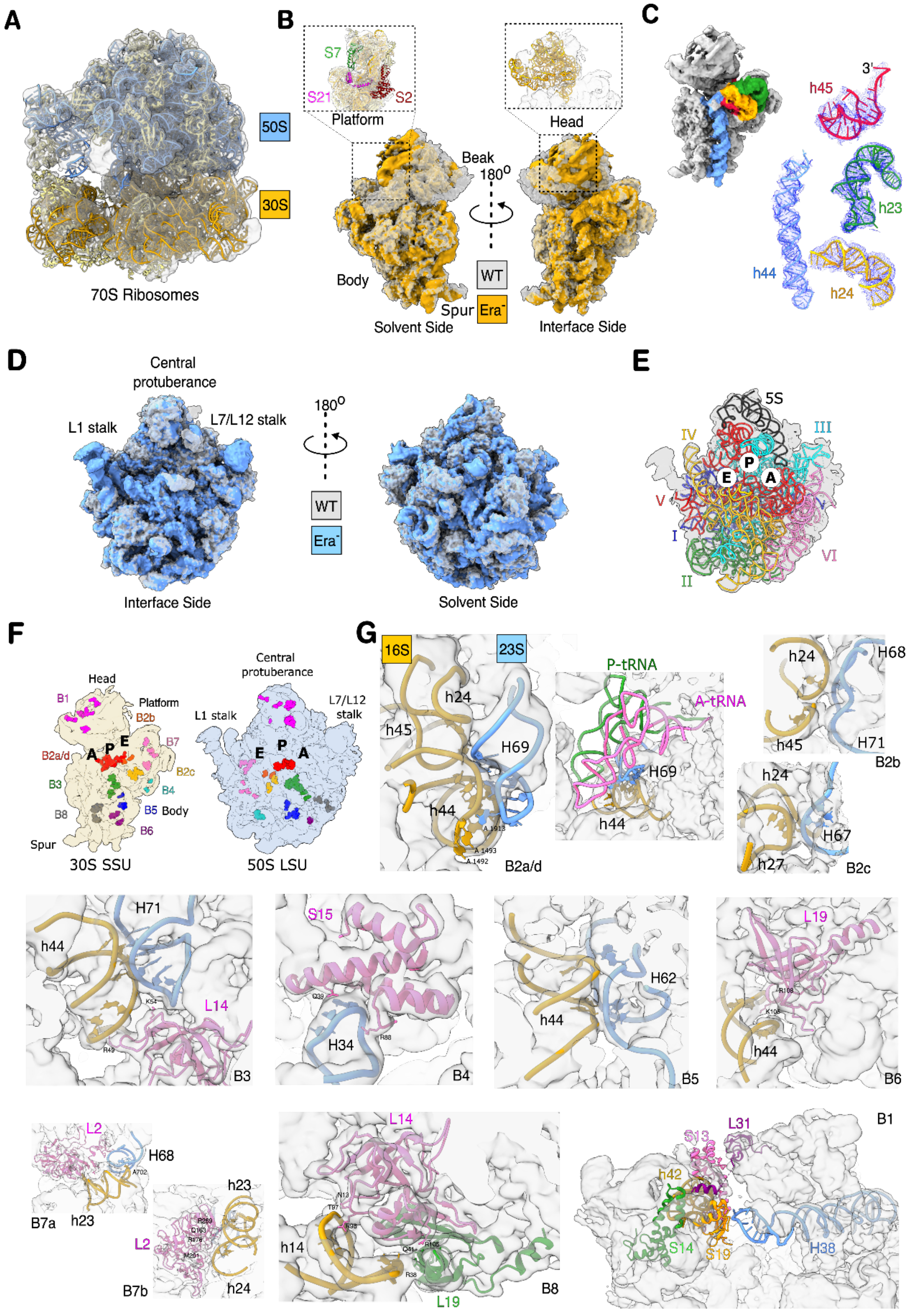

**Figure S5.** ***Analysis of ribosomes from Era-depleted cells***

**(A)** The ensemble Cryo-EM map of Era$^{-}$ 70S (grey transparent surface) fitted with atomic model (16S rRNA in yellow ribbon, 23S & 5S in blue ribbon and rProteins in khaki ribbon) of WT 70S (PDB ID: 7K00) is shown **(B)** Differences in WT model (grey surface) vs Era$^-$ 30S (yellow surface) from 70S map (A) shows the regions associated with the body are mature but the head (insert) and the rProteins in platform (insert) are disordered. **(C)** WT 16S rRNA model (ribbon) fitted with Era$^-$ 30S map (dark blue mesh) shows some of the important helices h23 (green), h24 (yellow), h44 (blue), h45 (red) associated with the decoding centre is folded into the correct conformation. **(D)** WT model of 50S (blue surface) fitted with Era$^-$ 50S Cryo-EM map (grey surface) shows no assembly defects. **(E)** WT model (cartoon) fitted with Era$^-$ 50S Cryo-EM map (surface in grey) shows the presence of various domains of 23S rRNA and 5S rRNA in their mature conformation. The P, A and E-site tRNA binding site on 50S are indicated. **(F)** Representative map indicating the position of intersubunit bridges on 30S and 50S subunits is shown. **(G)** The Era$^-$ 70S map (grey surface) fitted with WT model (23S rRNA helices in blue ribbon and 16S rRNA in yellow ribbon) shows that most of the 30S-50S intersubunit bridges are intact except for minor differences associated with B1 and B2a/d due to the flexible nature of the head.

---

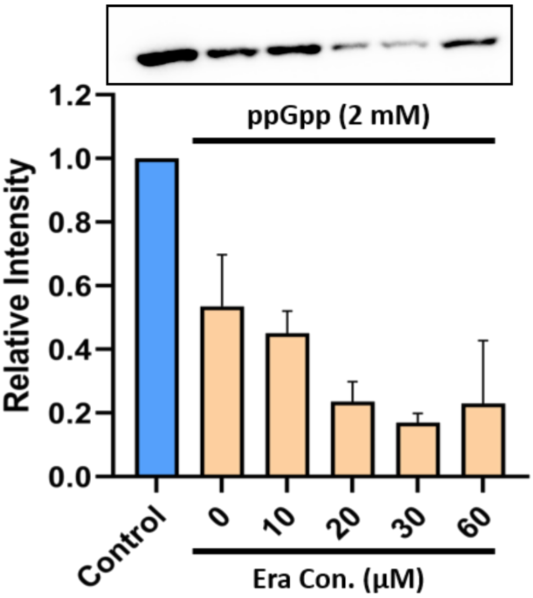

---

**Figure S6.** ***Effect of alarmones on translation***

*in vitro* translation assay showing that the addition of the alarmone ppGpp decreases GFP translation. However, it cannot be rescued by the incremental addition of Era as detected by Western blotting and densitometry. A representative blot of *in vitro* translation assay corresponding to the signal intensity for mEGFP translation is shown.

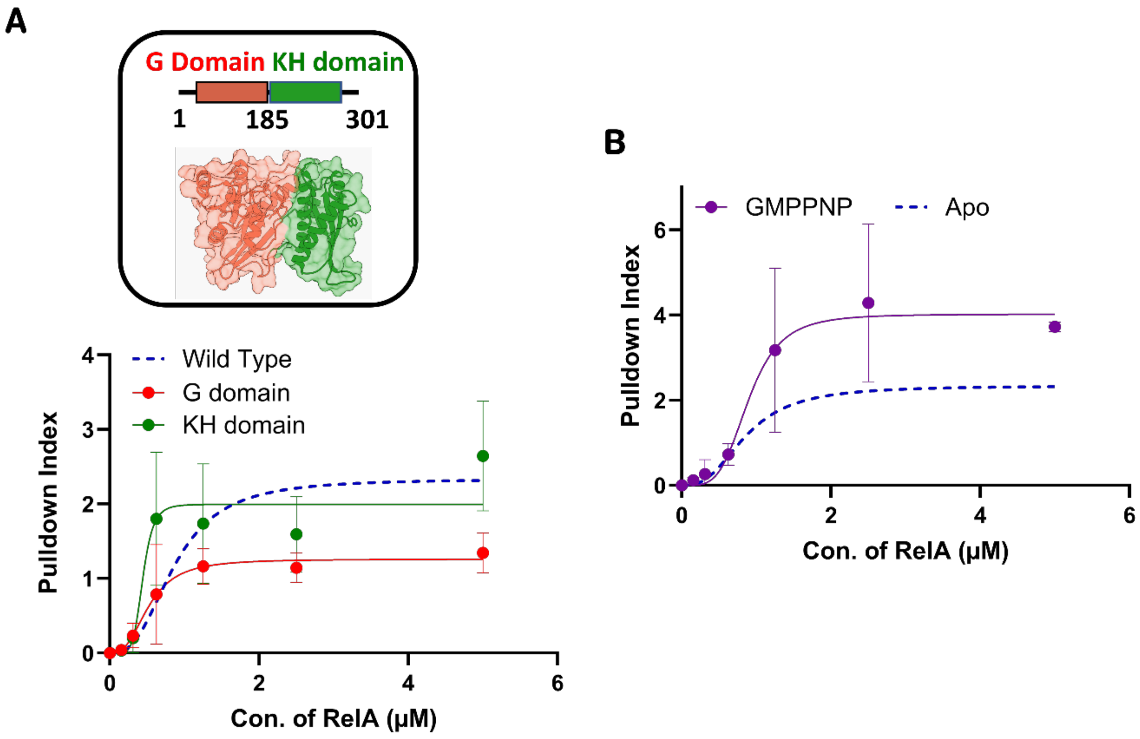

**Figure S7.** ***Interaction of G and KH-domain of Era with RelA***

**(A)** A model of Era (PDB ID: 1EGA) shows an N-terminal G-domain (red) and C-terminal KH-domain (green). Protein-protein pull-down assay shows both the G-domain and KH-domain of Era can independently interact with RelA, as analysed by Western blotting and densitometry. The representative binding curve of full-length Era is highlighted in a blue dotted line for reference. The G-domain (1-185 aa) and KH domain (186-301 aa) of Era interact with RelA with a $K_d$ of 0.523 μM ($n_H$ = 2.770, $R^2$ = 0.8148) and 0.441 μM ( $n_H$ = 6.248, $R^2$ = 0.735), respectively [n=3 biological replicates]. **(B)** Protein-protein pull-down assay shows interaction of RelA to Era in the presence of GMPPNP, a GTP analog ($K_d$ = 0.8966 µM, $n_H$ = 4.138, $R^2$ = 0.8037). The representative binding isotherm for Era in the apo state from (A) is shown in dotted blue line for reference [n=3 biological replicates].

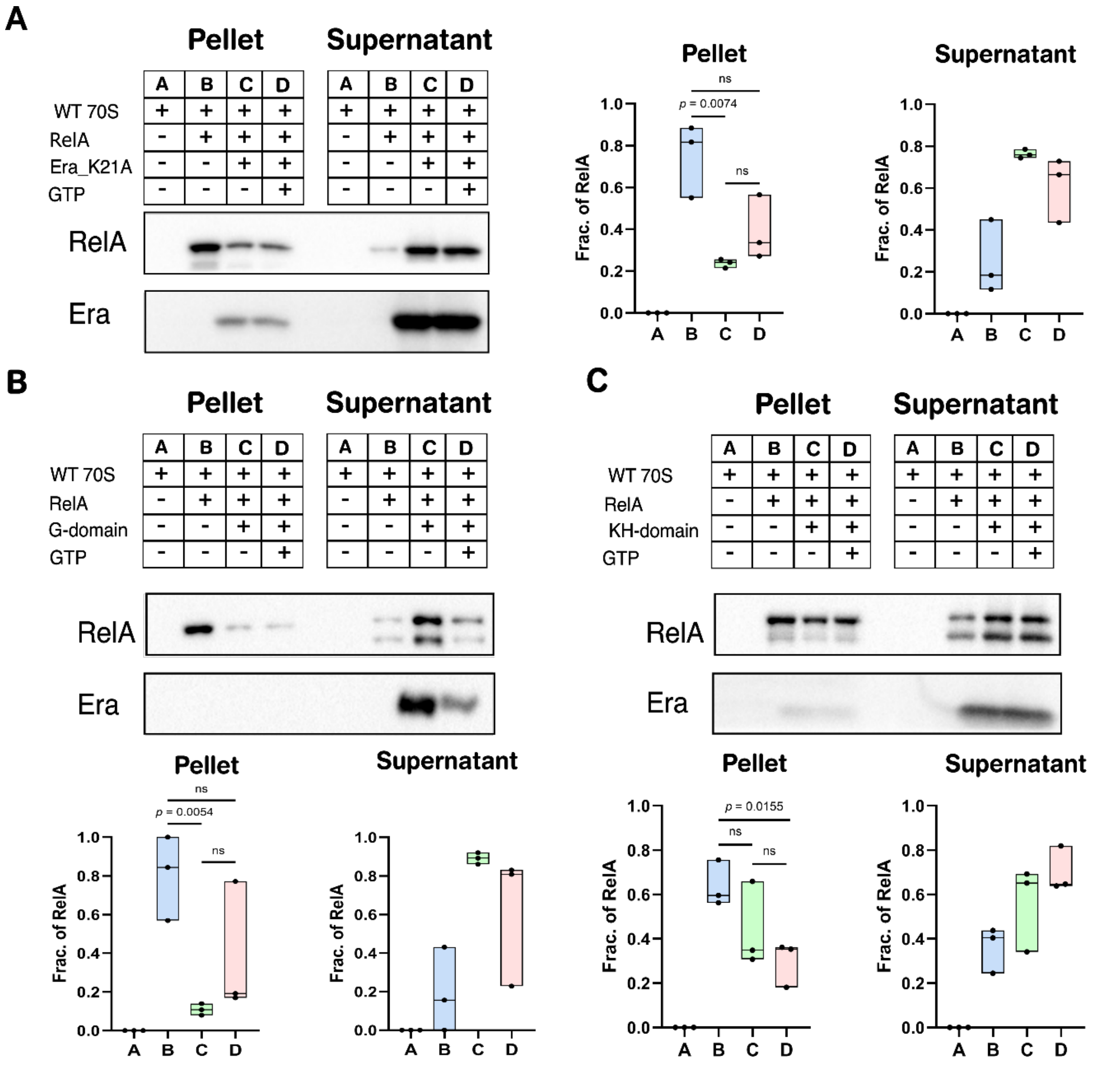

**Figure S8.** ***Effect of Era mutants on RelA and ribosome interaction***

**(A)** Co-sedimentation assay shows point mutant of Era (K21A), which lacks GTP binding, releases 70S-bound RelA. The release activity was analysed using Western blot and densitometry [Statistical analysis: Unpaired t-test, n=3 biological replicates]. **(B)** Co-sedimentation assay shows G-domain (1-185 aa) releases 70S-bound RelA. The release activity was analysed using western blot and densitometry [Statistical analysis: Unpaired t-test, n=3 biological replicates]. **(C)** Co-sedimentation assay shows KH-domain (186-301 aa) releases 70S-bound RelA, albeit with reduced activity in comparison to G-domain. The release activity was analysed using western blot and densitometry [Statistical analysis: Unpaired t-test, n=3 biological replicates]

### Table S1. List of strains and plasmids used in this study

| Strain/ Plasmids | Description | Source |
|---|---|---|
| BW25113 (Wt) (CGSC #7636) | *F-,Δ(araD-araB)567, ΔlacZ4787(::rrnB-3), λ-, rph-1, Δ(rhaD-rhaB)568, hsdR514* | *The E. coli Genetic Resource Center* (ECGRC) |
| TOP10 | *F- mcrA Δ(mrr-hsdRMS-mcrBC) ϕ80lacZΔM15 Δ lacX74 recA1 araD139 Δ(araleu)7697 galU, galK, rpsL endA1, nupG, StrR* | Invitrogen |
| B wild type (CGSC # 5365) | *lon-, malB-, dcm-* | *The E. coli Genetic Resource Center* (ECGRC) |
| K12*era*:HAssrA | *F-,Δ(araD-araB)567, ΔlacZ4787(::rrnB-3), λ-, rph-1, Δ(rhaD-rhaB)568, hsdR514, era*:HAssrA | This paper |
| K12*spoT*:HAssrA | *F-,Δ(araD-araB)567, ΔlacZ4787(::rrnB-3), λ-, rph-1, Δ(rhaD-rhaB)568, hsdR514, spoT*:HAssrA::kan | This paper |
| Δ*relA* | F-,Δ(araD-araB)567, ΔlacZ4787(::rrnB-3), λ-, rph-1, Δ(rhaD-rhaB)568, hsdR514, *era*:HAssrA, Δ*relA::smR* | This paper |
| pKD46 (CGSC#7739) | ori R101, repA101ts, AmpR, araC, expresses λ Red genes (gam-bet-exo) under the control of arabinose inducible promoter (pBAD). | *The E. coli Genetic Resource Center* (ECGRC) |
| BW25141/pKD13 | *F-, Δ(araD-araB)567, ΔlacZ4787(::rrnB-3), Δ(phoB-phoR)580, λ-, galU95, ΔuidA3::pir+, recA1, endA9(del-ins)::FRT, rph-1, Δ(rhaD-rhaB)568, hsdR514*, pKD13 | *The E. coli Genetic Resource Center* (ECGRC) |
| BW25141/pKD32 | *Genotype: F-, Δ(araD-araB)567, ΔlacZ4787(::rrnB-3), Δ(phoB-phoR)580, λ-, galU95, ΔuidA3::pir+, recA1, endA9(del-ins)::FRT, rph-1, Δ(rhaD-rhaB)568, hsdR514*, pKD32 | *The E. coli Genetic Resource Center* (ECGRC) |
| pET-1R | T7-lac inducible, ColE1 origin plasmid for expressing genes with N-terminally fused Strep-II tag. | Scott Gradia Addgene plasmid #29664 |
| pQE2 | ori ColE1, AmpR, expresses gene of interest to synthesise N-terminal 6xHis tagged protein under the IPTG inducible T5 promoter. | Qiagen |
| pET-13S-R | T7-lac inducible, ColE1 origin plasmid for expressing genes with N-terminally fused Strep-II tag. | Scott Gradia Addgene plasmid #48328 |

| | | |
|---|---|---|
| pEra-1R | T7-lac inducible, ColE1 origin plasmid for expressing WT *era* with N-terminally fused Strep-II tag. | This study |
| pHflX-1R | T7-lac inducible, ColE1 origin plasmid for expressing WT *hflX* with N-terminally fused Strep-II tag. | This study |
| pObgE-1R | T7-lac inducible, ColE1 origin plasmid for expressing WT *obgE* with N-terminally fused Strep-II tag. | This study |
| pEra-Gdomain-1R | T7-lac inducible, ColE1 origin plasmid for expressing only $era_{1\text{-}185}$ G-Domain with N-terminally fused Strep-II tag. | This study |
| pEra-KHdomain-1R | T7-lac inducible, ColE1 origin plasmid for expressing only $era_{186\text{-}301}$ KH-Domain with N-terminally fused Strep-II tag. | This study |
| pEraK21A-1R | T7-lac inducible, ColE1 origin plasmid for expressing only $era_{K21A}$ p-loop mutant with N-terminally fused Strep-II tag. | This study |
| pLJSRSF7 | T7-lac inducible, RSP 1030 origin plasmid for expressing gene with N-terminally fused 6His_MBP_SUMO tag. | Hideo Iwai Addgene #64693 |
| pLJ_RelA | T7-lac inducible, RSP 1030 origin plasmid for expressing only *relA* with N-terminally fused 6His_MBP_SUMO tag. | This study |
| pET His6 GFP TEV LIC vector (1GFP) | ori pMB1, KanR, lacI, expresses an enhanced version of gfp under the control of an IPTG inducible promoter (PT7lac). | Scott Gradia , Addgene plasmid #29663 |
| pmEGFP | ori ColE1, AmpR, expresses mEGFP with 6xHis tagged under the IPTG inducible T5 promoter. | This study |
| pT5mEGFP | pET-IR backbone, KanR, expresses mEGFP with 6xHis tagged under the IPTG inducible T5 promoter. | This study |
| pAUG-bgal | pQE2 carrying a lacz gene with a AUG start codon under the IPTG inducible T5 promoter. | Sharma et al. *Nucleic Acids Res.* 2019 |
| pT5AUG-bgal | pET-IR backbone, KanR, expresses AUG-bgal under the IPTG inducible T5 promoter. | This study |
| pBAD33-mf-lon | pBR322 origin vector expressing *Mesoplasma florum* lon protease under a pBAD promoter | Robert Sauer, Addgene plasmid #21867 |

### Table S2. List of oligonucleotides used in this study

| Oligo name | Sequence (5' to 3') | Description |
|---|---|---|
| gyrAqPCRFP | GAAGGCGATAAAGTCGTCTC | qPCR probe for *gyrA* |
| gyrAqPCRRP | ATCTGGTCGCAGTCATCTA | qPCR probe for *gyrA* |

| | | |
|---|---|---|
| cysGqPCRFP | CTGGTTGCTCTGCCTATTC | qPCR probe for *cysG* |
| cysGqPCRRP | CTGGCATTCCGTGTTCAA | qPCR probe for *cysG* |
| eraqPCRFP | ATCGAACGTTTCGTCTCTAAC | qPCR probe for *Era* |
| eraqPCRRP | CCCAACCGGATTTCACTTT | qPCR probe for *Era* |
| relAqPCRFP | CGTTCGCCGGATGTTATT | qPCR probe for *relA* |
| relAqPCRRP | GCAGTAATCTTCCAGTTCCC | qPCR probe for *relA* |
| spoTqPCRFP | GGACTTGGTAACGCAATGA | qPCR probe for *spoT* |
| spoTqPCRRP | GTGGATCACCAGACCTTTAC | qPCR probe for *spoT* |
| EraWT_1R_FP | CGCCGAAAACCTGTACTTCCAATCCAATATTATGAGCATCGATAAAAGTTA | Cloning *era* in to pET-1R vector |
| EraWT_1R_RP | GCTCGAATTCGGATCCGTTATCCACTTCCTTAAAGATCGTCAACGTAACC | Cloning *era* in to pET-1R vector |
| HflX_1R_FP | GCCGAAAACCTGTACTTCCAATCCAATATTTTGTTTGACCGTTATGATGC | Cloning *hflX* into pET-1R vector |
| HflX_1R_RP | GCTTGTCGACGGCGCTCGAATTCGGATCCTCATTAGATCAGGTAATCGATCA | Cloning *hflX* into pET-1R vector |
| ObgE_1R_FP | GCCGAAAACCTGTACTTCCAATCCAATATTATGAAGTTTGTTGATGAAGC | Cloning *obgE* into pET-1R vector |
| ObgE_1R_RP | GCTTGTCGACGGCGCTCGAATTCGGATCCTCATTAACGCTTGTAAATGAACT | Cloning *obgE* into pET-1R vector |
| Era1-185_Gdomain_1R_FP | GCCGAAAACCTGTACTTCCAATCCAATATTATGAGCATCGATAAAAGTTA | Cloning $era_{1\text{-}185}$ G-domain into pET-1R vector |

| | | |
|---|---|---|
| Era1-185_Gdomain_1R_RP | GCTCGAATTCGGATCCGTTATCCACTTCCTTATCAGATGTAATCTTCCGGGAAGT | Cloning *era*$_{1-185}$ G-domain into pET-1R vector |
| Era186-301_KHDomain_1R_FP | GCCGAAAACCTGTACTTCCAATCCAATATTACCGATCGCTCACAGCGTTT | Cloning *era*$_{186-301}$ KH-domain into pET-1R vector |
| Era186-301_KHDomain_1R_RP | CGCAAGCTTGTCGACGGCGCTCGAATTCGGATCCTTATCAAAGATCGTCAACGTA | Cloning *era*$_{186-301}$ KH-domain into pET-1R vector |
| EraK21A_1R_RP | TTTGTTCAACAATGTGGATGCGCCAACGTTCGGACGTCCG | Cloning *era*$_{K21A}$ into pET-1R vector |
| RelA_pLJSRSF7_FP | AGACTCACAGAGAACAGATTGGTGGATCCGGTGGAGGTATGGTTGCGGTAAGAAGTGC | Cloning *relA* into pLJSRSF7 vector |
| RelA_pLJSRSF7_RP | ACCAGACTCGAGTGCGGCCGCAAGCTTTTACTAACTCCCGTGCAACCGAC | Cloning *relA* into pLJSRSF7 vector |
| pQE2_mEgfp_FP | TAACTATGAAACATCACCATCACCATCACCATATGGTGAGCAAGGGCGAGGAGCT | Cloning *mrgfp* into pQE2 vector |
| pQE2_mEgfp_RP | ACAGGAGTCCAAGCTCAGCTAATTAAGCTTTCATTACTTGTACAGCTCGTCCA | Cloning *mrgfp* into pQE2 vector |
| ΔRelA_SmR_FP | TAGTTGCGATTTGCCGATTTCGGCAGGTCTGGTCCCTAAAGGAGAGGACGCGACCGAGTGAGCTAGCTAT | Used for creating knock-outs of *relA* |
| ΔRelA_SmR_RP | GTAGATACAGTATATATCAATCTACATTGTAGATACGAGCAAATTTCGGCGGCTTGAACGAATTGTTAGA | Used for creating knock-outs of *relA* |
| RelA200FLank_FP | ATAGTTTATGTATCCTGTAACCCTG | Gene flanking primers used for screening of *relA* null mutants |
| RelA200Flank_RP | TTTGCCATCCACCAGGTCAATCTTC | Gene flanking primers used for screening of *relA* null mutants |

| | | |
|---|---|---|
| SpoT_HA_FP | GCAAAATCCGCGTGATGCCAGACGTGATTAAAGTC ACCCGAAACCGAAATTATCCGTATGATGTTCCGG | Used for creating knock-in of *spoT* with HA tag at c-terminus |
| SpoT_HA_FP | CTGGCGAGCATTTCGCAGATGCGTGCATAACGTGT TGGGTTCATAAAACAACGGGCCCGGGATCCGATT | Used for creating knock-in of *spoT* with HA tag at c-terminus |
| SpoT200Flank_FP | TAGGCGGAAAGGATCCGCTG | Gene flanking primers used for screening of *spoT_HA* mutants |
| SpoT200Flank_RP | GCCGCTGCCGAAGCCATGGT | Gene flanking primers used for screening of *spoT_HA* mutants |
| Era200Flank_FP | TTGCAAGAATATTTGCAGGG | Gene flanking primers used for screening of *spoT_HA_ssrA* mutants |
| Era200Flank_RP | TGCTACGTTTTGGCGGGCGT | Gene flanking primers used for screening of *spoT_HA_ssrA* mutants |
| FP1.era-HA | GGGCCGACGACGAACGCGCACTGCGCAGTCTCGGT TACGTTGACGATCTTGAACAAAAAC | Used for creating knock-in of *era* with HA_SSRA tag at c-terminus |
| RP1.HA-ssra | GTTCGCCGCTCCAACCAGATCTTCTTCAGAAATAA GTTTTTGTTCAAGATCGTCA | |
| FP2_SSRA | TCTGAAGAAGATCTGGTTGGAGCGGCGAACAAAA AC | |
| RP2_SSRA | TAGATGCATTCGCGAGGTACTTACAACTGAAGAC GGTTCGCC | |
| FP3 | CGAACCGTCTTCAGTTGTAAGTACCTCGCGAATGC ATCTAGATT | |

| | |
|---|---|
| RP3 | GCGACTATGCAGGACAAATGCGCGCTGCCAGCCTT CCATCGGAGTTACTCACGGGCCCGGGATCCGATT |

### Table S3. List of reagents/enzymes/antibodies used in this study

| Reagent/enzymes/antibodies Name | Source | Catalog number |
|---|---|---|
| Rabbit anti-GFP primary antibody | Bio Bharati Life Science | BB-AB0065 |
| Goat anti-rabbit HRP conjugated secondary antibody | Invitrogen | A16096 |
| Mouse anti-HA Tag monoclonal primary antibody | Invitrogen | 26183 |
| Mouse anti-RelA monoclonal primary antibody | Santa Cruz Biotechnology | sc-81622 |
| Goat anti-mouse HRP conjugated secondary antibody | Invitrogen | A16066 |
| Mouse anti-GAPDH monoclonal primary antibody | Invitrogen | MA5-15738 |
| Mouse anti-Strep II-Tag monoclonal primary antibody | ABclonal technology | AE066 |
| SspI-HF® | New England Biolabs | R3132S |
| BamHI-HF® | New England Biolabs | R3136S |
| NdeI | New England Biolabs | R0111S |
| HindIII-HF® | New England Biolabs | R3104S |
| XhoI | New England Biolabs | R0146S |
| NheI-HF® | New England Biolabs | R3131S |
| DNase I (RNase-free) | New England Biolabs | M0303S |
| SuperSciptTM III Reverse Transcriptase | Invitrogen | 18080044 |
| SUPERase·In™ RNase Inhibitor | Invitrogen | AM2696 |
| PURExpress® Δ Ribosome Kit | New England Biolabs | E3313S |
| iTaq™ Universal SYBR® Green Supermix | Bio-RAD | 1725121 |
| Magne® Protein G | Promega | G7471 |
| Qubit Protein Broad Range Assay | Invitrogen | A50668 |
| 10X RT random primers | Applied Biosystems | 4319973 |

| | | |
|---|---|---|
| Qubit RNA BR Assay kit | Invitrogen | Q10210 |
| Clarity™ Western ECL Substrate | Bio-RAD | 1705060 |
| CelLytic™ B Cell Lysis Reagent | Sigma-Aldrich | C8740 |
| Lysozyme | HiMedia | MB098 |
| Isopropyl-b-D-thiogalactopyranoside | HiMedia | RM2578 |
| 2-Nitrophenyl β-D-galactopyranoside | Sigma-Aldrich | N1127 |
| d-Desthiobiotin | Sigma-Aldrich | D1411 |
| Trizol Reagent | Invitrogen | 15596018 |
| Phenol solution | Sigma-Aldrich | P4682 |
| Acridine orange | HiMedia | MB116 |
| Propidium iodide | HiMedia | TC252 |
| Sucrose | SRL | 27580 |
| Guanosine-3',5'-bisdiphosphate | Jena Biosciences | NU-884S |
| Guanosine-3',5'-pentaphosphate | Jena Biosciences | NU-885S |
| Guanosine 5′-triphosphate sodium salt hydrate | Sigma-Aldrich | G8877 |
| Guanosine 5′-diphosphate sodium salt | Sigma-Aldrich | G7127 |
| Guanosine 5′-[β,γ-imido]triphosphate trisodium salt hydrate | Sigma-Aldrich | G0635 |
| Dimethyl sulfoxide | Sigma-Aldrich | 472301 |
| MTT, 3-(4,5-Dimethyl-2-thiazolyl)-2,5-diphenyl-2H-tetrazolium Bromide | Sigma-Aldrich | 475989 |
| Chloramphenicol | Sigma-Aldrich | C0378 |
| Kanamycin sulphate | HiMedia | MB105 |
| Spectinomycin dihydrochloride pentahydrate | HiMedia | CMS6235 |
| Acetone | Sigma-Aldrich | 179124 |
| Methanol | Actylis | 67-56-1 |
| Tween 20 | MERCK | 9005-64-5 |
| Trichloroacetic acid | MERCK | 1.00807 |
| Chloroform | MERCK | 1.94506 |

| | | |
|---|---|---|
| Glycerol | FINAR | 56-81-5 |
| Primers | Integrated DNA Technologies (IDT) | |
| Precision Plus Protein™ Dual Color Standards | Bio-RAD | 1610374 |
| Phenylmethanesulfonyl fluoride | Sigma-Aldrich | P7626 |
| Imidazole | HiMedia | MB019 |
| 2-Mercaptoethanol | HiMedia | MB041 |
| L-(+)-Arabinose | Sigma-Aldrich | A3256 |
